# Immune Isoform Atlas: Landscape of alternative splicing in human immune cells

**DOI:** 10.1101/2022.09.13.507708

**Authors:** Jun Inamo, Akari Suzuki, Mahoko Ueda, Kensuke Yamaguchi, Hiroshi Nishida, Katsuya Suzuki, Yuko Kaneko, Tsutomu Takeuchi, Yasushi Ishihama, Kazuhiko Yamamoto, Yuta Kochi

## Abstract

Alternative splicing events are a major causal mechanism for complex traits, but they have been understudied due to the limitation of short-read sequencing. Here, we generated a comprehensive full-length isoform annotation of human immune cells, Immune Isoform Atlas, by long-read sequencing for 29 cell subsets. Our atlas contained a number of unannotated transcripts and isoforms such as a read-through transcript of *TOMM40-APOE*. We profiled functional characteristics of isoforms including encoded domains, inserted repetitive elements, and translational efficiency, and we showed that repetitive elements significantly explained the diversity of unannotated isoforms. Some of the isoforms are expressed in a cell-type specific manner, whose alternative 3’-UTRs usage contributed to their specificity. Further, we identified a number of disease-associated isoforms by isoform switch analysis and by integration of several quantitative trait loci analyses with genome-wide association study data. Our findings will promote the elucidation of the pathomechanism of diseases via alternative splicing.

## Introduction

Over 90% of human genes undergo alternative splicing, resulting in hundreds of thousands of transcript isoforms (Frankish et al., 2021; Pan et al., 2008; Wang et al., 2008). Alternative splicing can generate isoforms that differ in coding sequences through mechanisms that include exon skipping, a choice between mutually exclusive exons, the use of alternative splice sites, and intron retention, causing diversity in the open reading frame (ORF) and function of proteins (Baralle and Giudice, 2017; Xing and Lee, 2006). In addition, different 5′- and 3′-untranslated regions (UTRs) can quantitatively affect cellular functions of isoforms by altering translational efficiency and mRNA stability (Griesemer et al., 2021; Kim et al., 2021; Leppek et al., 2018). These splicing events can trigger human diseases, as genetic variants that affect alternative splicing, defined as splicing quantitative trait loci (sQTL), are enriched in the loci discovered by genome-wide association studies (GWAS) for complex diseases including immune-mediated diseases (IMDs) (GTEx Consortium, 2020; Lappalainen et al., 2013; Rotival et al., 2019). Indeed, in the GTEx project, 23% of the GWAS loci were co-localized with sQTL.

Generally, sQTL can be identified by testing the association of a variant’s genotype with the junction read counts of isoforms (Li et al., 2018), or alternatively, with the transcript-ratio of isoform expressions (Lappalainen et al., 2013). The latter method, also known as trQTL analysis, enables a direct understanding of which isoform expression is altered, that is, what kind of changes in the coding sequences or UTRs occur. However, the accuracy of quantification of junction reads and isoform expressions is susceptible to the credibility of isoform annotation. Therefore, accurate isoform annotation containing full-length sequences, even for those with low expression levels, can reveal novel disease pathogenetic mechanisms via alternative splicing. In fact, we have recently demonstrated that some low-expression isoforms that are disease-causing have coding sequences that were incomplete in the GENCODE annotation (Yamaguchi et al., 2022).

The emergence of long-read sequencing that can generate reads of 10,000 bases or more has revolutionized genomic studies; it has improved the mapping accuracy of reads, *de novo* assembly of genomes, and the detection of structural variations including repetitive elements (Amarasinghe et al., 2020; Sedlazeck et al., 2018). It has also advanced the precise evaluation of transcript structures as well as the discovery of novel transcripts that had been missed by short-read sequencing (Garalde et al., 2018; Gupta et al., 2018; Oikonomopoulos et al., 2016; Sharon et al., 2013; Tilgner et al., 2015; Weirather et al., 2017). Motivated by these features of long-read sequencing, large-scale projects aiming to reconstruct full-length transcripts using multi-tissue samples are ongoing (Glinos et al., 2022; Pardo-Palacios et al.). However, to our knowledge, there have been no studies focusing on immune cell subsets other than whole blood cells or lymphoblastoid cell-lines (LCL). Each of the diverse immune cell subsets has critical and specific functions in response to external stimuli (Lee et al., 2014; Quach et al., 2016; Schmiedel et al., 2018), the metabolic system (Lackey and Olefsky, 2016), and the nervous system (Ousman and Kubes, 2012), and their dysregulations are implicated in various human diseases (Fang et al., 2018). In addition, the importance of cell-type-specific alternative splicing in the immune system is known for a variety of genes, and cell-type-specific profiling of isoforms will help elucidate complicated immune system networks (Martinez and Lynch, 2013; Oberdoerffer et al., 2008; Ruan et al., 2011).

Here, we generated a comprehensive full-length isoform annotation, which we named the Immune Isoform Atlas, by sequencing 29 immune cell subsets using long-read sequencing. We also profiled the characteristics of isoforms among immune cell subsets, including transcription start sites (TSS), ORFs, and inserted transposable elements (TEs). We identified 2,575 isoforms expressed in a cell-type specific manner, whose 3′-UTR usage might mainly affect their cell-type-specific function. In addition, integrated analysis using the Immune Isoform Atlas and publicly available short-read RNA-seq datasets identified a significant number of isoforms that were switched between IMDs and controls. Furthermore, we also examined multiple kinds of QTL effects, including sQTL, on these isoforms, and we revealed disease-relevant isoforms by integrating these QTL with GWAS data. Our Immune Isoform Atlas (open on web; http://XXX) will bridge the gap between genomic and functional analysis and help elucidate the pathogenesis of IMDs.

## Results

### Overview of the Immune Isoform Atlas

To clarify the transcriptome profiles of immune cells at full-length levels, we isolated 29 immune cell subsets from the peripheral blood cells of a healthy donor. cDNA libraries were made from the poly(A) mRNA, PCR amplified, and subjected to long-read sequencing using the MinION Oxford Nanopore Technologies platform (**Methods; Figure 1A; Table S1-2**). The median number of raw reads in 29 subsets was 5,721,968 (**Figure S1A**). We mapped raw reads with quality > Q7 on the human genome (GRCh38) and filtered out reads with a mapping quality with MAPQ < 10, resulting in 4,566,622 reads as the median of 29 subsets (**Figure S1B**). After additional quality control using credible TSS annotation based on Cap Analysis of Gene Expression sequencing (CAGE-seq) datasets and removal of isoforms with redundant sequences (**Figure S1C**), we identified a total of 159,369 isoforms transcribed from 17,496 genomic loci, which we call the Immune Isoform Atlas. We compared our isoforms with those of GM12878 cell lines obtained by long-read sequencing (Workman et al., 2019), and 7.8% of our isoforms were matched.

**Figure 1.**
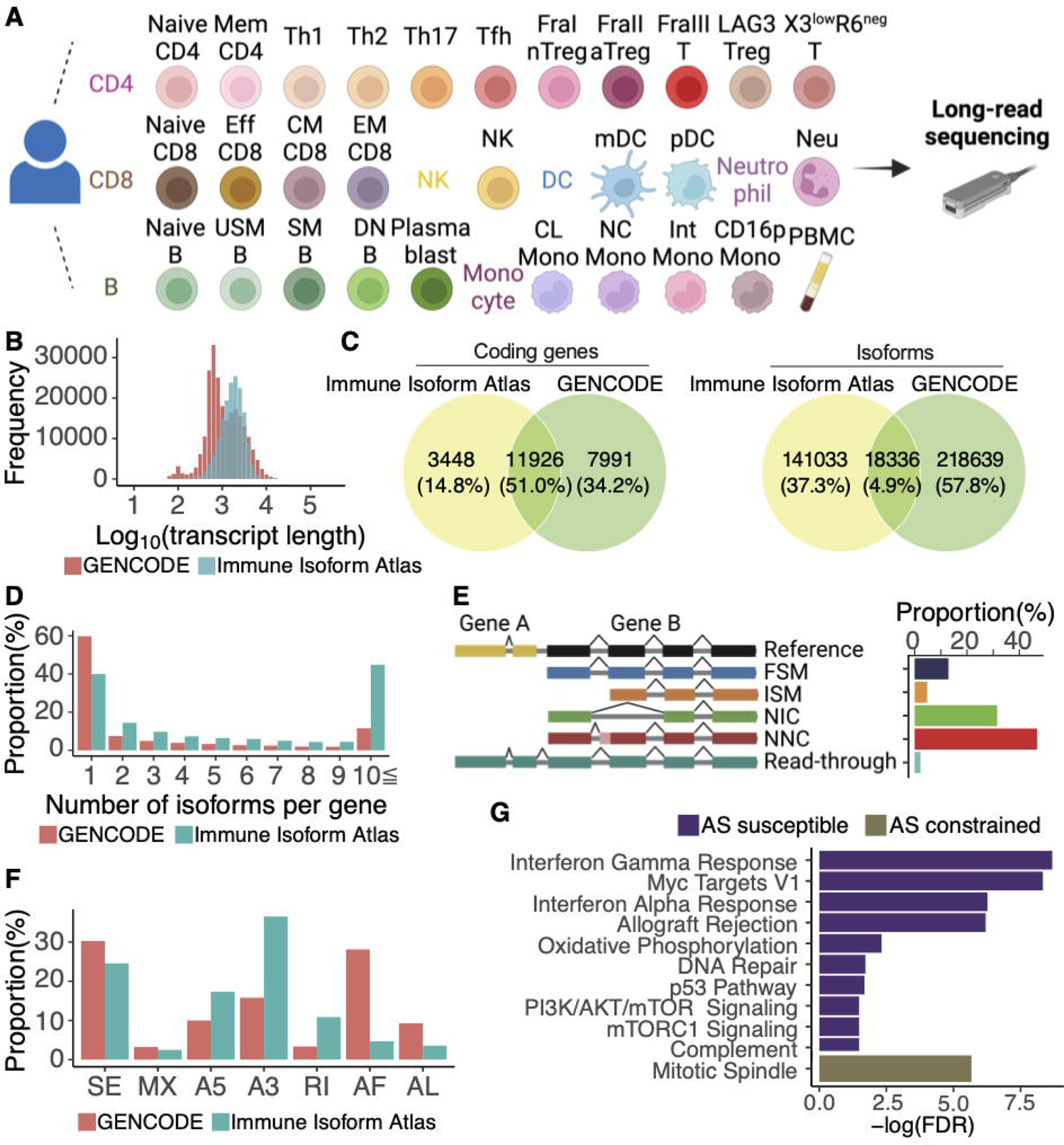
Overview of the Immune Isoform Atlas. (A) Summary of the cell subsets included in long-read sequencing in this study. A full description of the subset names and gating strategy is provided in **Table S1**. (B) The distribution of transcript length. (C) Overlap of coding genes (left) and isoforms (right) between the Immune Isoform Atlas and GENCODE. (D) The proportion of the number of alternatively spliced isoforms per genomic locus. (E) The proportion of structural categories of isoforms in the Immune Isoform Atlas. FSM (full splice match), meaning the reference and query isoform have the same number of exons, and each internal junction agrees; ISM (incomplete splice match), meaning the query isoform has fewer 5′ exons than the reference, but each internal junction agrees; NIC (novel in catalog), meaning the query isoform does not have an FSM or ISM match, but is using a combination of known donor/acceptor sites; NNC (novel not in catalog), meaning the query isoform does not have an FSM or ISM match, and has at least one donor or acceptor site that is not annotated. (F) The proportion of splicing events of isoforms in the Immune Isoform Atlas. SE, skipping exon; MX, mutually exclusive exon; A5, alternative 5′ splice site; A3, alternative 3′ splice site; RI, retained intron; AF, alternative first exon; AL, alternative last exon. (G) Pathway analysis using genes that have the top 5% (purple, alternative splicing (AS) susceptible genes) and bottom 5% (gold, AS constrained genes) of the number of alternatively spliced isoforms per gene after correction by the expression level.

The median length of isoforms in our atlas was 1,752 nucleotides (maximum 9,933 nucleotides), which was significantly longer than the isoforms registered in the comprehensive gene annotation of GENCODE version 38 (hereafter GENCODE) (median 929 nucleotides, maximum 347,561 nucleotides) (Wilcoxon test, p < 2.2e-16, **Figure 1B**). At the gene level, we found at least one transcript isoform in the Immune Isoform Atlas for 51% of the genomic loci where coding transcripts are annotated in GENCODE (**Figure 1C**). At the isoform level, we found 141,033 isoforms with sequences not registered in GENCODE (**Figure 1C**). The number of isoforms per genomic locus was higher in the Immune Isoform Atlas than in GENCODE, with 44% of the total genomic loci transcribing more than 10 different isoforms (**Figure 1D**). Comparing the percentage of transcriptional support level categories defined by GENCODE for the isoforms shared between GENCODE and the Immune Isoform Atlas, the most reliable category (all splice junctions are supported by at least one non-suspect mRNA) was the highest (**Figure S1D**). This supports the reliability of isoforms identified by long-read sequencing in our atlas.

We then examined what classes of alternative splicing occurred in our isoforms compared to those registered in GENCODE (**Figure 1E**). In view of splicing junctions, we found that 78% of the isoforms had either a novel splicing site (NNC) or a novel combination of known splice sites (NIC) as defined by SQANTI3 (Tardaguila et al., 2018). Regarding the type of splicing events using SUPPA2 (Trincado et al., 2018), intron retention, alternative 5′ splice sites, and alternative 3′ splice sites were more common in the Immune Isoform Atlas (**Figure 1F**). Since genes with higher expression levels had a greater number of alternatively spliced isoforms (**Figure S1E**), we corrected the number of alternative isoforms by the expression level of each gene. We then examined the characteristics of genes having the highest and lowest numbers of isoforms (top 5% and bottom 5%, respectively) (**Methods**). As a result, we found that genes involved in IFN signaling and mitotic spindles were enriched, respectively, suggesting that humans have acquired diverse isoforms related to immunological activity through alternative splicing, while cellular homeostasis was maintained by genes with a limited diversity of isoforms (**Figure 1G**).

### Novel coding transcripts predicted in the Immune Isoform Atlas

Next, we predicted the coding potential of isoforms by the GeneMark-ST algorithm (Tang et al., 2015), and identified 145,523 coding isoforms in total. To determine whether proteins are actually translated from the predicted ORFs, we referred to the data of ongoing proteome analysis using the LCL and a monocytic leukemia cell line (THP-1) (Nishida et al, manuscript in preparation). To reduce false positives and maximize the identification of novel proteins, we selected 16,190 isoforms expressed in LCL or THP-1 and did not exactly match the amino acid sequences registered in GENCODE. As a result, we confirmed peptides for 276 isoforms (**Table S3**), of which 150 peptides were not also registered in Swiss-Prot (UniProt Consortium, 2021).

Then, we focused on 3,448 genomic loci that differ from genes registered as coding sequences (CDS) in GENCODE (**Figure 1C**). Of note, 1,365 (40%) loci encoded so-called read-through isoforms, in which the mRNA extended through the conventional polyadenylation signal (PAS) but stopped at the PAS of adjacent genes or genomic loci (**Figure 2A**). Although read-through isoforms are known to be transcribed in particular situations in cells, such as in malignancy and infection (Grosso et al., 2015; Heinz et al., 2018), many read-through isoforms, such as *TOMM40_APOE* (**Figure 2B**), were also transcribed in cells under normal physiological conditions. Interestingly, the read-through isoform transcribed from the *TOMM40_APOE* locus was predicted to harbor conserved domains and conformational structure characteristics to both *TOMM40* and *APOE* (**Figure 2B-C**). In addition, 1,022 loci annotated as long non-coding RNAs (lncRNAs) in GENCODE were predicted as coding genes by GeneMark-ST (**Figure 2A**). Although lncRNAs are defined as over 200 nucleotides in length and do not code for a peptide or protein (Statello et al., 2021), previous studies have shown that a fraction of putative small ORFs within lncRNAs are translated (Ingolia et al., 2009; Slavoff et al., 2013). Notably, the genomic regions of predicted ORFs of these lncRNAs were more conserved compared to those of the UTRs and introns (**Figure 2D**), supporting their coding potential.

**Figure 2.**
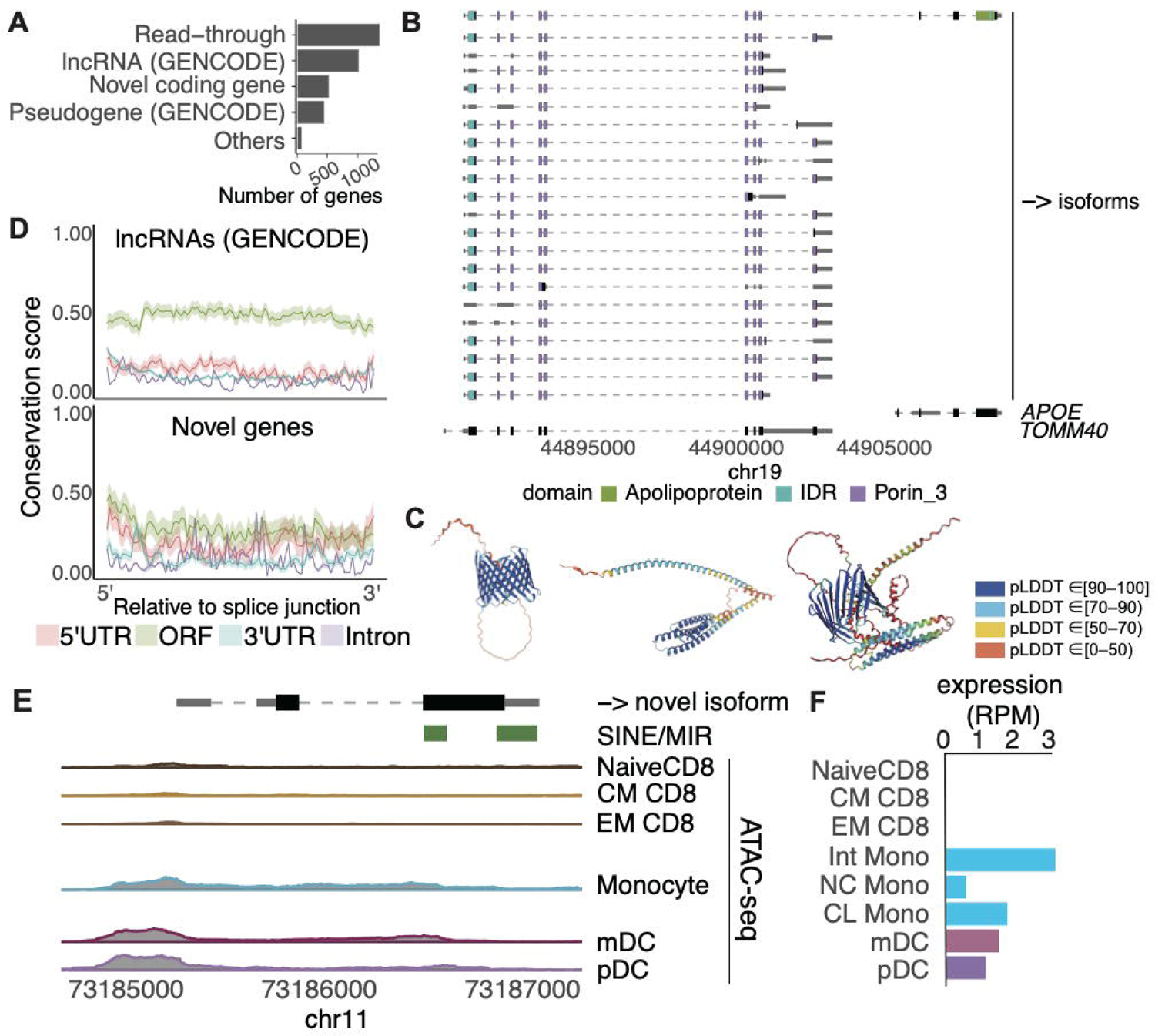
Novel coding genes identified in Immune Isoform Atlas. (A) The proportion of categories of transcripts encoded in genomic loci that differ from genes registered as CDS in GENCODE. (B) Example of a read-through isoform (top) transcribed from the *TOMM40* and *APOE* locus. The arrow indicates the direction of transcription in the genome. The collapsed gene structure registered in GENCODE is shown at the bottom. (C) Protein 3D structures of TOMM40 (left), APOE (center), and the read-though isoform (right) predicted using AlphaFold2 (Jumper et al., 2021). pLDDT is a per-residue estimate of its confidence on a scale from 0 - 100. Regions with pLDDT > 90, between 70 and 90, between 50 and 70, and < 50 are expected to be high accuracy, well (a generally good backbone prediction), low confidence, and a reasonably strong predictor of disorder, respectively. (D) Conservation score (Siepel et al., 2005) of predicted-coding transcripts registered as lncRNAs in GENCODE (top) and novel transcripts that were transcribed from genes not registered as coding in GENCODE (bottom) according to their region. Shading indicates ±1.96 standard deviation. (E) Example of an isoform from a novel gene locus. Peaks of ATAC-seq derived from relevant non-stimulated immune cell subsets around this locus are shown. The arrow indicates the direction of transcription in the genome. (F) The expression of the novel isoform in each cell subset is shown. The expression value was normalized by reads per million (RPM).

Next, we examined 529 loci encoding novel predicted coding genes, which did not overlap any annotated genomic loci in GENCODE (**Figure 2A**). Notably, their predicted ORFs were highly conserved (**Figure 2D**). We examined whether these transcripts were registered in the Comprehensive Human Expressed SequenceS (CHESS) database, which contains an inclusive set of genes based on nearly 10,000 RNA-seq experiments (Pertea et al., 2018). Most of the transcripts were found in CHESS, but the remaining 47 transcripts from mutually exclusive loci did not overlap any annotated genes in CHESS. For example, one novel transcript from chromosome 11 had a predicted 448 nucleotide ORF (**Figure 2E**), and notably, we found an open chromatin region from the TSS to intronic regions in subsets of monocytes and dendritic cells using datasets of the assay for transposase-accessible chromatin sequencing (ATAC-seq) (Calderon et al., 2019). This corresponded to our expression profile, in which it is only expressed in these subsets in the Immune Isoform Atlas (**Figure 2F**). Because the ORF and 3′-UTR of this isoform contained sequences derived from short interspersed elements (SINE)/MIR, the insertion of these TEs may have made it difficult to map the reads by short-read sequencing.

### Transposable elements inserted in isoforms

Motivated by the finding of the novel transcript above, we investigated the contribution of TEs in constituting novel genes (i.e., those not registered in GENCODE). TEs are major repetitive elements and make up approximately half of the human genome (Lander et al., 2001). TEs can be subdivided into four major categories: (i) DNA transposons; (ii) long terminal repeat (LTR) retrotransposons; (iii) long interspersed elements (LINEs); and (iv) SINEs. We comprehensively searched for repetitive elements including TEs from curated libraries of Dfam (Storer et al., 2021) and Repbase (Bao et al., 2015) that are inserted in the transcripts in our atlas. Interestingly, repetitive elements were inserted more in those from the novel genomic loci (83.9% vs. 47.6%, chi-square test, p < 0.001), indicating that the Immune Isoform Atlas included substantial transcripts missed by short-read sequencing due to the insertion of repetitive elements.

We also examined the contribution of TEs in the splicing diversity of known genes in the Immune Isoform Atlas. The median length of inserted repetitive elements including TEs was 188 nucleotides (**Figure 3A**). The maximum length was 3,675 nucleotides, which was a LINE/L1 inserted in the 3′-UTR of a *CCDC7* isoform (**Figure S2**). The TEs in all isoforms were summed for each class, and the most common were SINEs (**Figure 3B**). Comparing the newly identified isoforms with those common to GENCODE, the former had more TEs inserted (70% vs. 52%, chi-square test, p < 0.001, **Figure 3C**). The distribution of TEs around gene bodies was non-random, with the highest number of insertions in the 3′-UTR and the lowest numbers in the TSS (**Figure 3D**). We then examined the enrichment of each TE class at each position of the isoforms by comparing their proportions with those of the entire genome. Interestingly, the proportion of TEs was non-uniform; LTRs, which are autonomous and coding TEs, were enriched in TSSs and ORFs, while SINEs were enriched in 5′-UTRs and 3′-UTRs (**Figure 3D**). Among the isoforms with TEs, that of *LGALS3*, which has numerous functions in the immune system (Liu and Rabinovich, 2010), was transcribed from an alternative TSS. Because LTR/ERVL-derived sequences were inserted around the TSS (**Figure 3E**), this LTR has brought an alternative promoter as well as an additional CDS into this gene. A region-by-region comparison of the conservation (phastCons score > 0.8) (Siepel et al., 2005) of genomic loci with and without repetitive element insertions showed that the inserted regions were significantly less conserved (TSS, odds ratio (OR) = 0.42 [95% confidence interval (95%CI): 0.32-0.53]; ORFstart, OR = 0.58 [95%CI: 0.50-0.66]; ORFend, OR = 0.60 [95%CI: 0.56-0.65]; TTS, OR = 0.49 [95%CI: 0.46-0.52]; splicing junction, OR = 0.71 [95%CI: 0.70-0.73]), indicating that repetitive elements were inserted into these regions after humans diverged from other mammals, and these contributed to the diversity of isoforms.

**Figure 3.**
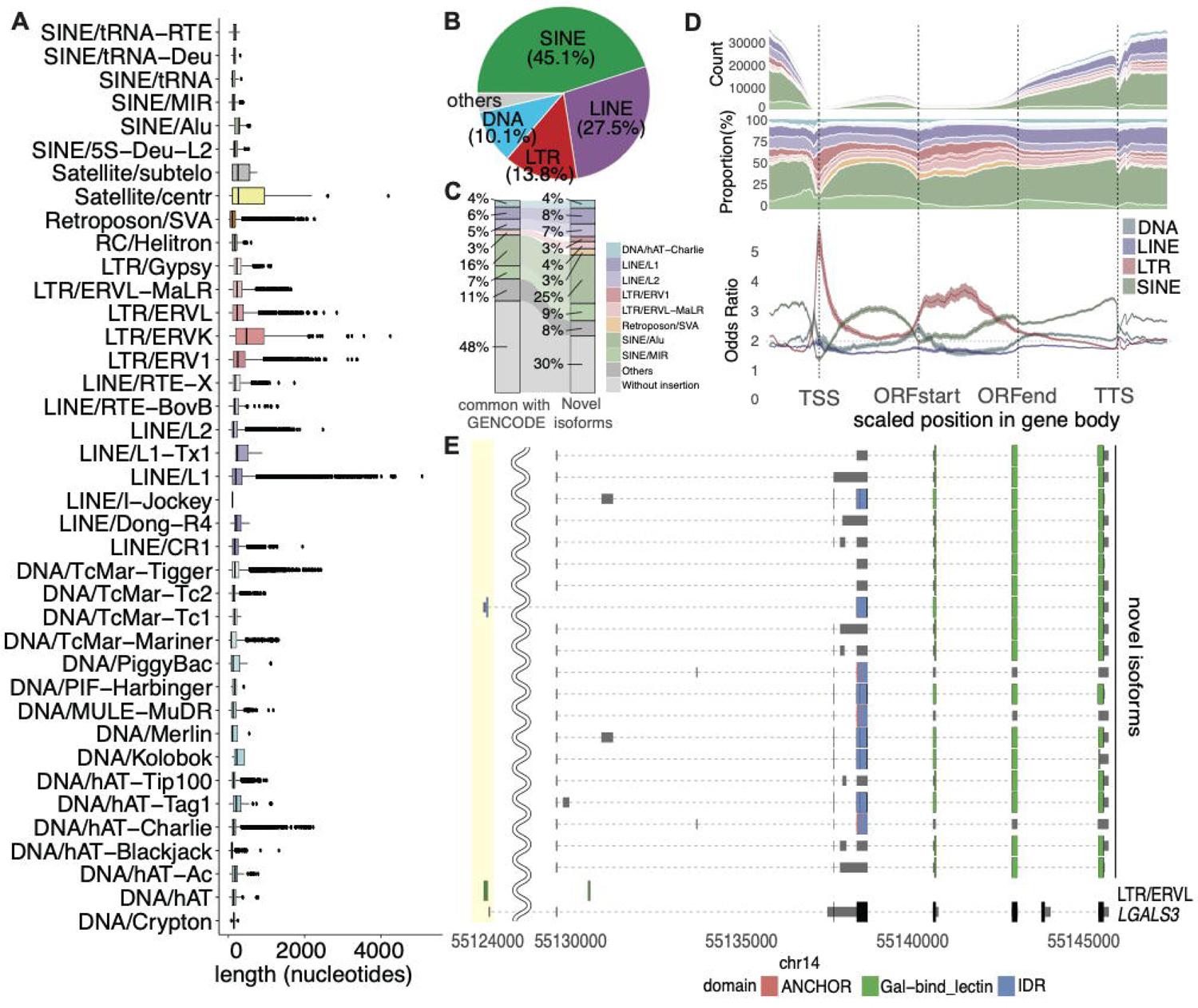
Repetitive elements inserted in isoforms. (A) The length of inserted repetitive elements in isoforms according to their class. The color of boxplot indicates their classes. (B) The proportion of class families of the total repetitive elements inserted in the isoforms. (C) Comparison of the proportion of class families of the total repetitive elements inserted in isoforms between novel isoforms and those that are also registered in GENCODE. (D) The distribution of inserted repetitive elements in gene bodies ± 1000 bp. The x-axis is a scaled position relative to the TSS. The y-axis is the counts (top), proportion (center), and the enrichment of each TE class family at the scaled position (bottom). The color represents the class of repetitive elements corresponding with **Figure 3A**. (E) Example of isoforms with an inserted LTR at the alternative TSS highlighted in yellow. The collapsed gene structure registered in GENCODE is shown at the bottom.

### Isoforms expressed in a cell-type-specific manner

Each immune cell subset expresses cell-type-specific genes, such as those encoding cytokines and transcriptional factors, involved in their respective cellular functions (Ota et al., 2021; Schmiedel et al., 2018). Because some of these are regulated at the isoform level (e.g., a spliced isoform of *RORG* is essential in Th17 cells (Ruan et al., 2011)), cluster analysis based on the isoform ratio, that is the ratio of isoform abundance over the total gene abundance, should connect subsets with identical lineages. To test this hypothesis, we first performed an unsupervised hierarchical clustering analysis based on the similarity of the isoform ratio in each related gene. As expected, we found that subsets of an identical lineage (e.g., B-cell subsets) were in close proximity to each other, suggesting that the abundance of isoforms may reflect the functional characteristic of each cell type (**Figure 4A**). Then, we examined the isoforms expressed in a cell-specific manner utilizing Shannon entropy as an index of cell specificity and identified 2,575 isoforms for 29 cell subsets (**Figure 4B; Table S4**) (Kadota et al., 2006; Schug et al., 2005). This cell-type specificity was recapitulated in the expression data of the relevant immune cell subsets obtained from short-read RNA-seq (Schmiedel et al., 2018) (**Figure S3A-B**).

**Figure 4.**
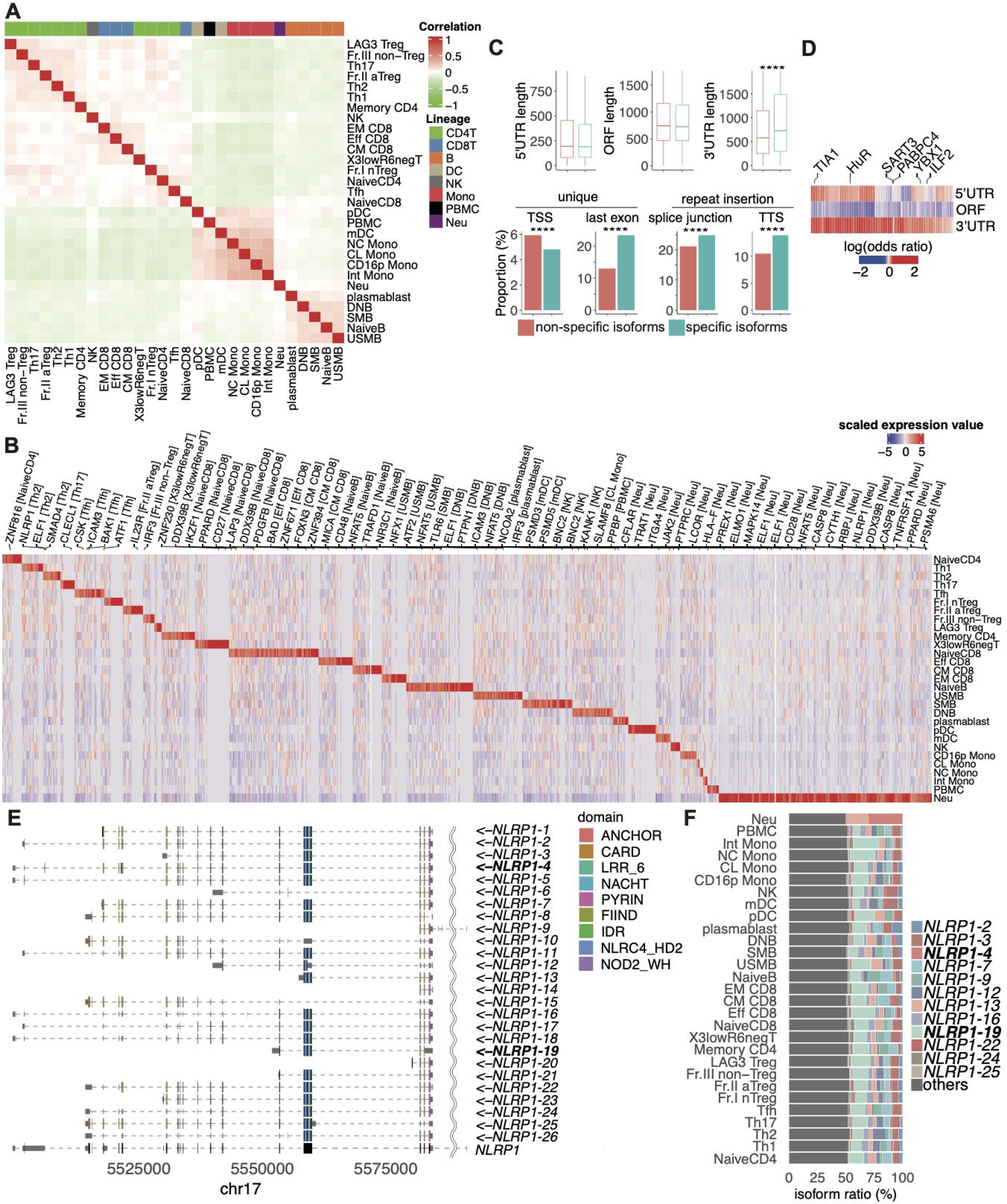
Cell-type-specific isoforms and their characteristics. (A) Unsupervised hierarchical clustering based on isoform ratios that were the top 5000 large expression variances among cell types. The colored bars indicate the cell lineages. (B) Cell-type-specific expressed isoforms. Column-wise Z scores of normalized counts are plotted. Representative gene symbols related to specific isoforms are annotated at the top. Full lists of cell-type-specific isoforms are provided in **Table S4**. (C) The length of the 5′-UTR, ORF, and 3′-UTR (top) and the proportion of unique TSS, last exon, insertion of repetitive elements in splicing junctions, and TTS (bottom) compared between cell-type-specific isoforms and others. (D) Enrichment of each RBP motif according to gene regions compared between cell-type-specific isoforms and others. Representative significant RBPs (FDR < 0.01) are annotated at the top. (E) Example of a specifically expressed isoform transcribed from the *NLRP1* locus. The collapsed gene structure registered in GENCODE is shown at the bottom. (F) Isoform ratio in *NLRP1*. Arrows indicate the direction of transcription in the genome. The significance of comparison is as follows: ****, p<0.0001; ***, p<0.001; **, p<0.01; *, p<0.05. Each box reflects the interquartile range (IQR), and the whiskers of the box plots reflect the maximum and minimum values within each grouping no further than 1.5× IQR from the hinge.

As cell-type-specific expression of isoforms may occur at the transcriptional level (e.g., alternative TSS usage) and at the post-transcriptional level (e.g., alternative usage of splicing sites or PAS) (Trapnell et al., 2010, 2012), we examined which mechanism is prominent in the cell-type-specific isoforms. We found that while the proportion of unique TSSs was lower in the cell-type-specific isoforms compared with others (4.8% vs. 5.9%, chi-square test, p = 0.019, **Figure 4C**), these isoforms have longer 3′-UTRs (729 nucleotides vs. 579 nucleotides, Wilcoxon test, p < 0.001) and a higher proportion of unique sequences of the last exon (23.4% vs. 13.0%, chi-square test, p < 0.001), suggesting that the cell-type-specific isoforms prefer the usage of alternative splicing sites at the 3′-UTR and PAS. Interestingly, the insertion rates of repetitive elements into junction sites (24.9% vs. 21.1%, chi-square test, p < 0.001, **Figure 4C**) and TTSs (13.9% vs. 10.5%, chi-square test, p < 0.001) was higher in cell-type-specific isoforms. In addition, the proportion of non-canonical PAS (16.7% vs. 13.6%, chi-square test, p < 0.001) was higher in cell-type-specific isoforms as well. Motif enrichment analysis revealed that *GCCTGG* was the most enriched motif in the non-canonical PAS sequences.

As RBP binding is a major mechanism of alternative splicing (Fu and Ares, 2014), we hypothesized that RBP contributes to post-transcriptional regulation of cell-type-specific expression. We comprehensively searched for RBP binding motifs and compared them between cell-type-specific isoforms and others. As a result, many kinds of RBP binding motifs were enriched in 3′-UTRs (**Figure 4D**).

We show examples of cell-type-specific isoforms in **Figure 4E**. The isoform ratio of *NLRP1*-4 was the highest in neutrophils, while the isoform ratio of *NLRP1*-19, which lacks the FIIND domain essential for *NLRP1* inflammasome activity (Finger et al., 2012), was dominant in other cell types. In addition, the isoform ratio of *IL23R*, which essential for the differentiation of Th17 cells (McGeachy et al., 2009), was distinct in Th17 subset in comparison with other cell types (**Figure S3C-D**). Notably, while a cell-type specific isoform (*IL23R-5*) was identified in Th17 cells, another isoform (*IL23R-2*) specific for activated-Treg cells was also identified. Because the CDSs of these isoforms were different, their differential expression may contribute to Th17 and Treg functions and their plasticity (Kleinewietfeld and Hafler, 2013).

### Regulation of translation efficiency by isoform sequences

The efficiency of protein synthesis is governed by the regulatory elements in the 5′-UTR, ORF, and 3′-UTR (Jia et al., 2020; Leppek et al., 2018; Presnyak et al., 2015). As a classic example, a strong Kozak sequence immediately before the first codon improves start codon recognition as a feature of highly translated mRNAs (Jia et al., 2020). In addition, the secondary structure of mRNA may block or recruit ribosomes and other regulatory factors to enable a rapid, dynamic response to diverse cellular conditions (Leppek et al., 2018). Therefore, we speculated that examining the translational efficiency of each transcript in the Immune Isoform Atlas would provide additional insights into the regulation of translation and bridge the knowledge gap between transcripts and proteins. For this purpose, we remapped the Ribo-seq and RNA-seq data to the Immune Isoform Atlas, respectively, and calculated the scores of translational efficiency at the isoform level (**Methods; Figure 5A**). As expected, isoforms with a strong Kozak context score had higher translation efficiency scores (**Figure 5B**). Interestingly, the association between translational efficiency and the lengths of the 5- and 3-UTRs was reversed; i.e., shorter 5′-UTRs, as well as longer 3′-UTRs, were associated with higher translation efficiency (**Figure 5B**). Then, we calculated the correlation between the local folding strength of RNA and the scores of translation efficiency (**Methods**). We confirmed that the local folding strengths calculated for the sequences of each isoform were significantly correlated with the Parallel Analysis of RNA Structure (PARS) score, which was calculated from two high-throughput sequencing libraries per sample and provided profiling of the secondary structure at a single nucleotide resolution (Kertesz et al., 2010; Wan et al., 2014) (R = 0.67, p < 2.2e-16, **Methods; Figure S4**), indicating the credibility of local folding strengths. In addition, we found a negative correlation between the local folding strength of the 5′ leader (5′ end of an isoform) and higher translation efficiency (**Figure 5C**). In contrast, the local folding strength immediately after the first codon was positively correlated with the translational efficiency. These results are consistent with previous findings: stable RNA secondary structures at the 5′ leader, such as cap-proximal hairpins, block the assembly of the 43S pre-initiation complex onto the 5-UTR, while a hairpin positioned downstream of a first codon enhances translational initiation (Kozak, 1986, 1990), warranting our analysis of translation efficiency at the isoform level. As for the 3′-UTR, there was one scaled position with a significant negative correlation, which may reflect the complex association of RBP binding, miRNAs, and secondary structure (Kim et al., 2021).

**Figure 5.**
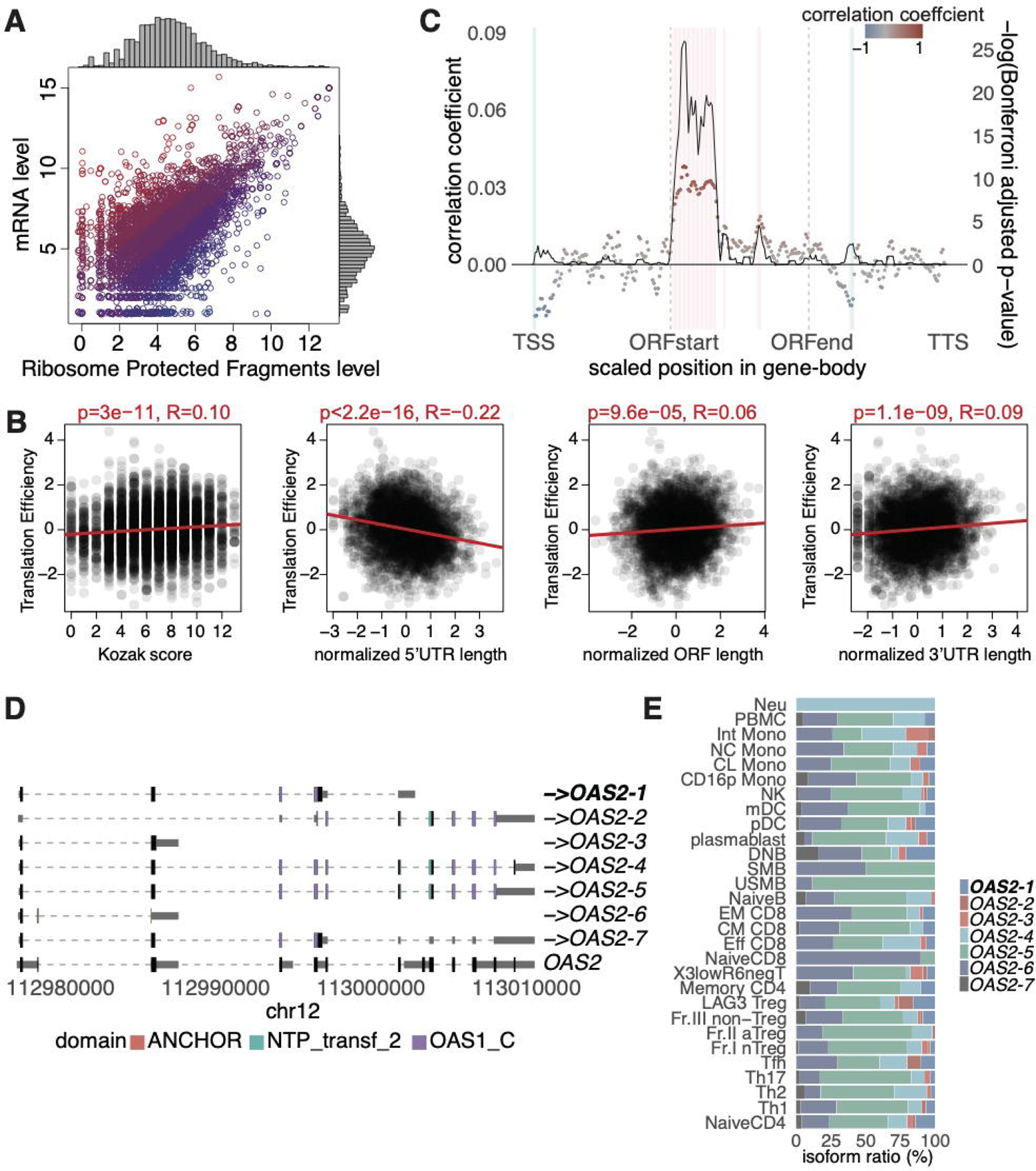
Translational efficiency at the isoform level. (A) Scatter plot of isoforms with normalized read counts of RNA-seq and Ribo-seq. Each dot represents a particular isoform and is colored by the rank of translational efficiency, and blue and red indicate high and low translational efficiency, respectively. (B) Correlation plots of translational efficiency scores with Kozak context scores (left), length of 5′-UTR (left center), ORF (right center), and 3′-UTR (right). (C) Correlation between translational efficiency and local folding strength of isoforms. The x-axis is a scaled position relative to the TSS. The solid line and colored points represent -log_10_ (Bonferroni adjusted p-value) and correlation coefficient, respectively. The areas with colored rectangles were significantly (Bonferroni adjusted p-value < 0.05) correlated at the scaled position in the gene body. (D) Example of the effectively translated isoform transcribed from the *OAS2* locus. The collapsed gene structure registered in GENCODE is shown at the bottom. The effectively translated isoform (top 10%) is highlighted in bold. (E) Isoform ratio in *OAS2*.

Furthermore, we investigated whether other features unique to the isoforms were associated with translational efficiency. For that purpose, we compared the characteristics of isoforms having the highest and lowest scores of translational efficiency (top 10% and bottom 10%, respectively). As a result, we found that having a unique TSS was associated with higher translation efficiency (7.0% vs. 2.3%, chi-square test, p < 0.001). In addition, those encoding a non-canonical ORF, which was defined as the first codon other than methionine using GeneMarkS-T (Tang et al., 2015), were associated with higher translation efficiency (26.1% vs. 17.6%, chi-square test, p < 0.001; **Table S5**). Non-normal ORFs can be translated into proteins and have received much attention in immunity and cancers (Jackson et al., 2018; Prensner et al., 2021). In the Immune Isoform Atlas, 28,837 isoforms were predicted to encode non-canonical ORFs. Of these, we examined 4,780 isoforms for peptides by proteome analysis, and found that 22 isoforms translated peptides from the non-canonical ORFs (**Table S3**). A specific example of isoforms encoding non-canonical ORF in our atlas is the *LGALS3*-isoform, already noted above (**Figure 3E**). This isoform has a unique TSS and non-canonical ORF with serine predicted as the first codon, and its translation efficiency was in the top 10%. Another example of effectively translated isoforms was *OAS2-1* (**Figure 5D**). This isoform has the shortest 5′-UTR and longest 3′-UTR among the isoforms transcribed from the *OAS2* locus. The translational efficiency of *OAS2-1* was in the top 10% and expressed specifically in double negative (DNB) B cells (**Figure 5E**).

### Isoforms abundantly spliced in IMDs

To investigate whether newly identified isoforms are involved in disease pathogenesis, we compared the abundance of isoforms between case and control subjects, taking systemic lupus erythematosus (SLE) as an example. SLE is an autoimmune disease with activation of interferon signature genes known to be involved, though details of the mechanism at the isoform level are not well understood (Tsokos et al., 2016). To investigate pathogenic isoforms in SLE, we remapped short-read RNA-seq datasets obtained from whole blood cells of SLE and healthy subjects to the Immune Isoform Atlas (SLE = 99, healthy subjects = 18) (Hung et al., 2015) (**Methods**). As a result, we identified 84 genes whose isoform fractions were significantly switched between SLE and healthy individuals (false discovery rate (FDR) < 0.05). Among them, IRAK1 transduces signals from TLR7 and TLR9 by phosphorylating IRF7 to promote IFNα transcription (Tun-Kyi et al., 2011). One known isoform of *IRAK1* (*ENST 00000393687.6*) contains a protein kinase domain, but the novel *IRAK1-1* lacks this domain (**Figure 6A; Figure S5**). Although there was no difference in gene-level expression between case and control samples **(Figure 6B)**, the isoform fraction significantly switched, resulting in higher expression of a functional isoform (*ENST 00000393687.6*) in SLE **(Figure 6C)**. This implied that TLR7/9-IRAK1-IRF7 pathway activation due to upregulated expression of the functional *IRAK1* isoform (*ENST 00000393687.6*) in SLE may contribute to type 1 IFN dysregulation.

**Figure 6.**
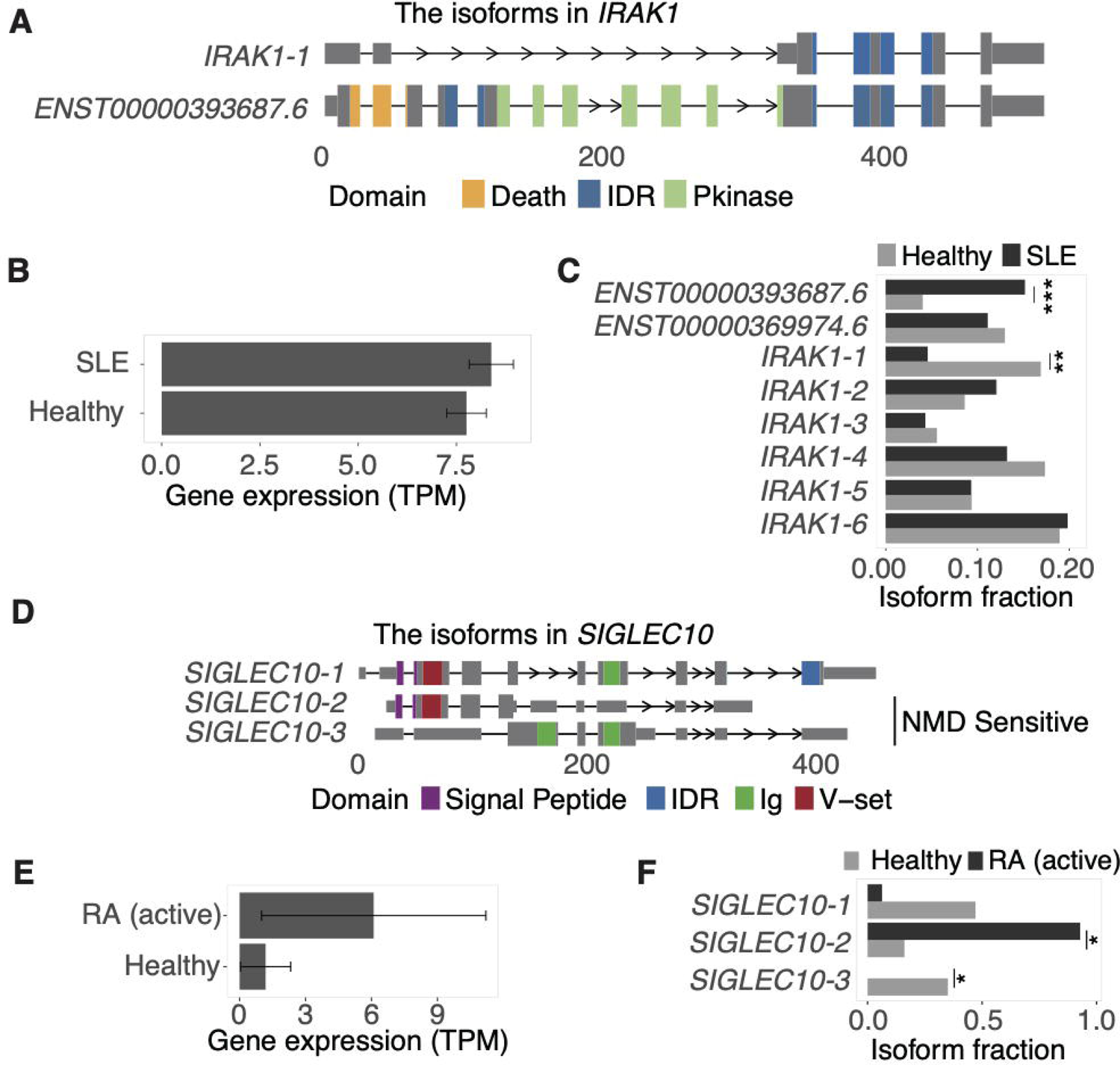
Switched isoforms in IMDs. (A) Structures of isoforms that are transcribed from the *IRAK1* locus and switched between SLE and healthy controls. The x-axis shows distance from the TSS. (B) Expression of *IRAK1* at the gene level. (C) Isoform fractions in *IRAK1*. (D) Structures of isoforms that are transcribed from the *SIGLEC10* locus and switched between RA (active) and healthy controls. (E) Expression of *SIGLEC10* at the gene level. (F) Isoform fractions in *SIGLEC10*. The gene expression value was normalized by scaledTPM, which scaled abundance estimates up to library size (Soneson et al., 2015). Error bars indicate ± 1.96 standard deviations. The significance of comparison is as follows: ***, p<0.001; **, p<0.01; *, p<0.05.

Rheumatoid arthritis (RA) is another example of an autoimmune disease characterized by chronic inflammation of the synovial tissue (McInnes and Schett, 2011). We applied our atlas to a single immune subset, CD45RA-positive effector memory (Temra) CD8^+^ T cells (active RA = 9, healthy subjects = 9) (Inamo et al., 2021; Takeshita et al., 2019). We found that three novel isoforms in the *SIGLEC10* locus, which suppresses inflammatory responses to damage-associated molecular patterns by interacting with CD24 (Chen et al., 2014), were differentially expressed in Temra CD8^+^ T cells from active RA and healthy controls subjects (**Figure 6D**). Two of them (novel isoforms 2 and 3) were predicted as being sensitive to nonsense-mediated mRNA decay (NMD), which is a surveillance pathway that degrades RNA and prevents the production of abnormal proteins (Lykke-Andersen and Jensen, 2015). At the gene level, *SIGLEC10* was more highly expressed in RA in comparison with healthy individuals (**Figure 6E**). At the isoform level, however, NMD-sensitive isoforms were dominant in RA (**Figure 6F**). This suggests that even though the expression was increased at the gene level, the relative expression of aberrant isoforms that are susceptible to NMD was increased, resulting in a relatively low inflammatory suppressive function of *SIGLEC10* in RA.

### QTL analysis and integrated analysis with GWAS

Analysis of differentially expressed isoforms as presented above identified disease-relevant isoforms, but they may simply reflect the disease course (i.e., they result from the disease). If alternative splicing is regulated by genetic variants that are defined as sQTL, and meanwhile the variants present susceptibility to disease, the variants and the splicing events should be causal to the disease. Thus, integration of sQTL analysis and GWAS data can comprehensively reveal isoforms involved in disease pathogenesis.

We first examined the impact of different reference annotations on junction-based sQTL. We remapped RNA-seq data of LCL samples derived from European subjects in the GEUVADIS cohort (Lappalainen et al., 2013) with each of two annotations, GENCODE and the Immune Isoform Atlas, independently, and compared the number of junctions with significant sQTL (FDR < 0.05). Of the total, 15% were identified only in the Immune Isoform Atlas as a reference (**Figure 7A**). To further ensure that sQTLs identified only in our atlas were not simply false positives, we verified that sQTL variants were significantly enriched in the GWAS variants compared with genome-wide variants (**Figure 7B**).

**Figure 7.**
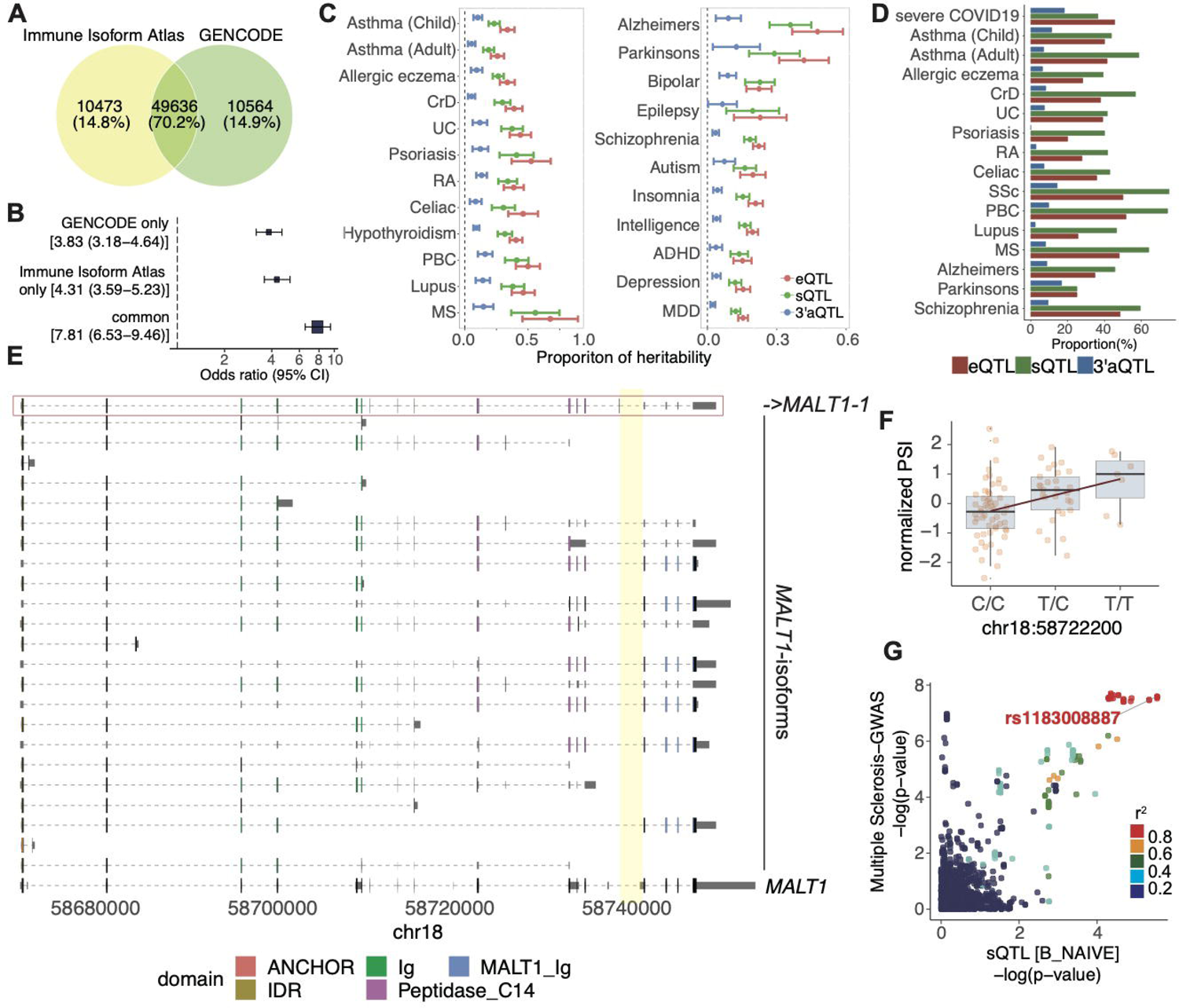
QTL and colocalization analysis with GWAS data. (A) Venn diagram for junctions with significant sQTL signals (FDR<0.05) in comparison with the Immune Isoform Atlas and GENCODE as a reference. (B) The enrichment of junction-based sQTL variants in GWAS variants compared to randomly selected 10,000 genome-wide variants (autosomes, minor allele frequency > 0.01 in European subjects of 1000 Genomes Phase 3 (Auton et al., 2015)). The list of GWAS data is available in **Table S6**. (C) The proportion of the heritability of QTL variants in IMDs (left) and neurological diseases (right). Error bar indicates ±1.96 standard deviation. CrD, Crohn’s Disease; UC, ulcerative colitis; RA, rheumatoid arthritis; PBC, primary biliary cirrhosis; MS, multiple sclerosis; ADHD, attention-deficit hyperactivity disorder; MDD, major depressive disorder. (D) The proportion of colocalized loci with each kind of QTL variants among all GWAS loci. SSc, Systemic Sclerosis. (E) Structures of isoforms transcribed from *MALT1* with a percent spliced in (PSI) value of a unique junction (chr18:58739121-58741865, highlighted in yellow) that is strongly associated with the GWAS signal of multiple sclerosis in naïve B cells. The related isoform to the unique junction (top) is framed by the red line. The collapsed gene structure registered in GENCODE is shown at the bottom. (F) QTL plot of normalized PSI of the unique junction in naïve B cells according to each genotype. (G) Colocalization plot of sQTL related and GWAS of multiple sclerosis.

Next, we remapped RNA-seq datasets from various cell conditions (Lappalainen et al., 2013; Lee et al., 2014; Quach et al., 2016; Schmiedel et al., 2018) to refine the mapping of sQTL so that we could identify pathogenic isoforms using the Immune Isoform Atlas. In addition to junction-based sQTL, we performed eQTL for genetic variants that affect gene expression, and 3′aQTL (Li et al., 2021), which associates cis-acting genetic variants with alternative polyadenylation (APA). The numbers of genes or isoform-related genes having effects on eQTL, sQTL, and 3′aQTL in one or more cell conditions were 9775, 10189, and 4087, respectively (nominally significant p-values < 1 × 10^-5^ (Aguet et al., 2017)). Genes involved in the IFN signaling pathway were enriched in 3′aQTL-eGenes, as previously reported (**Figure S6A**) (Li et al., 2021). As expected from a previous study (GTEx Consortium, 2020), the number of QTL variants increased with the sample size for each cell type (**Figure S6B**). eQTLs and 3′aQTL variants were abundant around TSS and TTS, respectively, while sQTLs were distributed throughout the gene body (**Figure S6C**). We then examined the proportion of each QTL in the heritability of IMDs and neurological diseases using stratified linkage disequilibrium score regression analyses (S-LDSCs) (Finucane et al., 2015), and found that many QTLs significantly contributed to the heritability of both disease types (**Figure 7C**).

Finally, we evaluated the colocalization between GWAS loci for IMDs and neurological diseases (16 GWAS data in total, **Table S6**) and QTL signals using coloc (PP4 > 0.8). To take advantage of the Immune Isoform Atlas, which contains full-length information on isoforms, and to detect more potential pathogenic isoforms, we also performed trQTL analysis. As a result, we found that 20–60% of GWAS loci colocalized with eQTL, sQTL (junction-based sQTL and trQTL), and up to 20% of GWAS loci with 3′aQTL (**Figure 7D; Table S7**). Notably, eQTL signal related to 63 genes of 3,448 genomic loci that were specific to the Immune Isoform Atlas (**Figure 1C**) had colocalization with any GWAS loci. Among the sQTL colocalized with GWAS loci, an SNP in *MALT1* (rs11873030), which was associated with multiple sclerosis (International Multiple Sclerosis Genetics Consortium et al., 2011), had an sQTL effect for a junction read (chr18:58739121-58741865) unique to *MALT1*-1; this isoform decreased with the risk allele (C allele) for the disease (**Figure 7E-G**). MALT1 transduces NF-kappaB (NFκB) signaling by antigen receptor stimulation, and importantly, the isoform with the sQTL effect lacks the MALT1 C-terminal immunoglobulin-like domain (**Figure 7E**).

## Discussion

Our datasets, obtained by long-read RNA sequencing of 29 immune cell subsets, provide a comprehensive full-length isoform profiling of the human immune system. We found an enormous complexity of splicing events that are substantially shaped by insertions of TEs into the human genome. This complexity brought by TEs may have been underestimated, owing to the difficulty in read-mapping of TEs obtained by short-read sequencing. However, our atlas shows that TEs provide isoform diversity to immunological genes by introducing alternative TSS, splicing sites, and polyadenylation signals, as has been shown in previous studies (Miao et al., 2020; Belancio et al., 2006; Roy-Engel et al., 2005). Considering their non-uniform distribution in the gene body, each class of TE has a unique role in adding functions to the immunological genes. These additional functions obtained by TEs could have been naturally selected by the environment, because immune cells play a critical role in protecting the host from external pathogens such as viruses or bacteria by triggering local and systemic inflammation. Indeed, a recent study demonstrated that the short isoform of the *ACE2* gene, which encodes a receptor for SARS-CoV-2, is upregulated by an interferon-inducible alternative promoter introduced by MIRb and LTR16A1 elements (Ng et al., 2020).

Our atlas contained many read-through transcripts, which are one of the most understudied categories of transcript isoforms, especially with regards to disease genetics. The *TOMM40_APOE* locus, where a novel read-through isoform was identified, is the most significant locus for Alzheimer’s disease (Jansen et al., 2019). Notably, trQTL signals of the read-through isoform in non-stimulated monocytes were marginally colocalized with GWAS signals (PP4 = 0.74, **Figure S6D-E**). The disease risk allele of rs204468 increased the isoform ratio of the read-through isoform among the isoforms transcribed from the *TOMM40* locus (trQTL beta = 0.64, p = 7.1 × 10^-6^). Because multiple independent association signals have been detected in this locus (Jansen et al., 2019), in addition to the well-established variant of *APOE4*, additional variants regulating the ratio of read-through isoform may also contribute to the pathogenesis of Alzheimer’s disease.

We also demonstrated cell-type-specific alternative splicing in the immune cell subsets. This may elucidate the role of genes in each immune cell subset that regulates inflammatory signals, as exemplified by the isoform of *NLRP1*. As a major mechanism for cell-type-specific expression, we found that alternative usage of the 3′-UTR was important, and it may affect the cellular function of isoforms, especially translational efficiency. We found that the length of the 5′-UTR and 3′-UTR had different effects; longer 3′-UTRs displayed higher translational efficiency. The length of the 3-UTR has immensely increased during eukaryotic evolution in comparison with the 5′-UTR (Pesole et al., 2001), indicating that they may have acquired different contributions to cellular functions. Interestingly, we also found that isoforms encoding non-canonical ORFs were translated efficiently, as confirmed by proteomic analysis. Although distinct roles of proteins translated by non-canonical ORF have been demonstrated in specific contexts, including gene regulation (Wright et al., 2022) and immune responses (Jackson et al., 2018), we should further examine whether these novel proteins that arise from non-canonical ORFs identified by our atlas have differential functions compared to those from conventional ORFs.

Our sQTL analysis showed that the Immune Isoform Atlas, used as a reference annotation, can identify functional junction-based sQTL variants that were missed when using GENCODE. Since existing alignment tools (Dobin et al., 2013; Kim et al., 2013) preferred to align junction reads to known junctions, the use of annotations having more complete and accurate information on the splicing junctions is critical for sQTL analysis. Furthermore, our atlas directly provides full-length sequences for isoforms identified in trQTL analysis as well as those in sQTL analysis that have unique junction sequences.

Finally, we identified novel pathogenic mechanisms via alternative splicing by performing isoform switch analysis and integrated analysis of QTLs and GWAS data. Gene-level analyses, such as differentially expressed gene analysis and eQTL analysis, alone cannot capture changes in the coding sequences and functional regions of alternatively spliced isoforms, which sometimes leads to misunderstanding of a disease as in the case of *SIGLEC10.* Our data highlight the necessity of alternative splicing analysis with annotation of full-length sequences. Use of short-read RNA-seq datasets and the Immune Isoform Atlas will facilitate the discovery of unknown pathogeneses of diseases and new therapeutic targets.

### Limitations of the study

There are several drawbacks to this study. First, the cell sample used was from a single healthy individual. Although it has been shown that differences between individuals are smaller than differences between cell types in alternative splicing (Wang et al., 2008), adding additional subjects for profiling isoforms by long-read sequencing will identify novel isoforms. Second, the use of bulk-derived CAGE-seq datasets in the process of filtering reliable isoforms may have excluded those with cell-type-specific TSS with subtle CAGE-peaks. Third, we did not condition the QTL analyses by other QTL effects. Different kinds of QTL signals are often detected in the context of a strong linkage disequilibrium structure, and it is sometimes difficult to detect independent effects. Last, the findings of predicted functional changes in spliced isoforms are supported by disease genetic studies and other omics studies (e.g., GWAS etc.), but experimental validation should be obtained in future studies.

## Acknowledgments

Computations were partially performed on the NIG supercomputer at the ROIS National Institute of Genetics. This study was supported by the Japan Society for the Promotion of Science (JSPS) Grant-in-Aids for JSPS Fellows (grant numbers: 21J00596 and 21J15131), Grant-in-Aids for Scientific Research (B) (grant numbers: 18H02849, 21H02459, and 22H02597), Grant-in-Aid for Scientific Research (S) (grant numbers: 18043381), and Grant-in-Aid for Challenging Research (grant number: 21K19501) from the MEXT of Japan, and grants from Nanken-Kyoten, TMDU and Medical Research Center Initiative for High Depth Omics. We thank K. Kobayashi for her technical assistance.

## Author contribution

J.I. conducted bioinformatics analysis with the help of M.U., K.Yamaguchi., and Y.Kochi, A.S managed and contributed to sample collection, cell sorting, and long-read RNA sequencing. H.N and Y.I. contributed to proteome analysis. K. Yamamoto, and Y.Kochi designed and managed the project. K.S., Y. Kaneko and T.T. provided the short-read RNA sequencing datasets for isoform switch analysis. J.I. and Y.Kochi wrote the manuscript with contributions from all authors to the final version of the manuscript.

## Declaration of interest

The authors declare no competing interests.

## Lead contact

Further information and requests for resources should be directed to and will be fulfilled by the Lead Contact, Yuta Kochi.

## Materials availability

This study did not generate new unique reagents.

## Data and code availability

Isoform expression data were deposited in the National Bioscience Database Center (NBDC) Human Database (https://humandbs.biosciencedbc.jp/en/) with the accession number XXXXXXXX. We used publicly available software for the analyses. Source code is available at https://github.com/juninamo/isoform_atlas.

## METHOD DETAILS

### Sample collections

We sorted PBMCs into 29 immune cell subsets from a healthy volunteer (42-year-old female) using a 14-color cell sorter, BD FACSAria Fusion (BD Biosciences), with purity > 99% using a MoFlo XDP instrument (Beckman Coulter) (**Table S1; Table S6**). Erythrocytes were lysed with potassium ammonium chloride buffer, and non-specific binding was blocked with Fc-gamma receptor antibodies. Sorted cells were lysed and stored at −80°C. Total RNA was extracted using the MagMAX total RNA kit (Ambion, Life Technologies). Total RNA preparation (100 μg) was added to 100 μL nuclease-free water and poly-A selected using NEXTflex Poly(A) Beads (BIOO Scientific) according to the manufacturer’s instructions and stored at −80°C. We intended to collect 5,000 cells with at least 1,000 cells per subset (5,000 cells were collected for > 80% of samples). We followed previously reported immune cell definitions provided by the Human Immunology Project (Maecker et al., 2012) for the flow cytometry staining panel with slight modification due to the availability of labeled antibodies. In addition, CXCR3^low^CCR6^-^ (X3^low^R6^neg^) T cells and LAG3^+^ Treg cells were sorted following previous studies, respectively (Nikitina et al., 2018; Okamura et al., 2015). Neutrophils were collected with EasySep Direct Human Neutrophil Isolation Kits (STEMCELL Technologies) or MACSxpress Neutrophil Isolation Kits human (Miltenyi Biotec) with an aim of 2 × 10^6^ cells, lysed, and stored at −80°C, followed by RNA isolation with an RNeasy Mini Kit (QIAGEN). This study was approved by the Ethics Committees of the Medical Research Institute, Tokyo Medical and Dental University and RIKEN Center for Integrative Medical Sciences. Written informed consent was obtained from the volunteer. Age and sex of the participant are deposited in the NBDC with the same accession number as denoted above.

### Long-read RNA sequencing and processing

We prepared cDNA libraries with a SMART-seq v4 Ultra Low Input RNA Kit (Takara Bio) and SQK-LSK109 (Oxford Nanopore Technologies). We used SMARTScribe Reverse Transcriptase for cDNA synthesis and SeqAmp DNA Polymerase for PCR amplification (20 cycles) of cDNA. A hundred fmol of cDNAs were sequenced by MinION Flow Cell (R9.4.1, FLO-MIN116; Oxford Nanopore Technologies) for 48 hours. Basecalling was performed using Guppy with the SUP (super high accuracy) model. We aligned reads to the GRCh38 genome reference using minimap2 (Li, 2018) with default parameters and reference to the splice junctions in the GENCODE annotation. We used SAMtools (Li et al., 2009b) to filter out reads with a mapping quality (MAPQ) less than 10. Then, we used the flair pipeline (Tang et al., 2020) to identify the full-length of isoforms and filtered them using the following criteria: 1) transcripts whose 5′ end was located within 100 bp from the TSS annotated by refTSS (Abugessaisa et al., 2019) and/or TSSclassifier (“relaxed” or “strict”), which are based on the FANTOM CAGE (Cap Analysis of Gene Expression) peak (Forrest et al., 2014), were extracted, 2) transcripts that have at least three full-length supporting reads (80% coverage and spanning 25 bp of the first and last exons) in total with the “--stringent” option. For splicing junction correction in the process of the flair pipeline, we used realigned reads with the GENCODE annotation obtained from 30 short-read RNA-seq datasets of immune cells (Lappalainen et al., 2013; Lee et al., 2014; Quach et al., 2016; Schmiedel et al., 2018).

Then, we used SQANTI3 with default parameters to filter out transcripts that are considered artifacts by intra-priming and reverse-transcription switching (Tardaguila et al., 2018).

Transcripts were compared to Workman et al. flair-called transcripts (Workman et al., 2019) using gffcompare (Pertea and Pertea, 2020) with the “--strict-match” option, which only allows a limited variation of the outer coordinates of the terminal exons by at most 100 bases.

### Genes susceptible to alternative splicing

To investigate the characteristics of genes that are susceptible to alternative splicing, we performed linear regression with the objective variable as the number of alternatively spliced isoforms and the explanatory variable as the read counts of genes in long-read sequencing. Then, we extracted genes with the top 5% highest and lowest 5% residuals by the *resid* function in the stats R package and performed pathway analysis using Enrichr (Chen et al., 2013) independently.

### ATAC-seq

ATAC-seq data of immune cells (Calderon et al., 2019) were obtained from GEO under accession GSE118189. As in the original paper, we processed raw reads as follows: we trimmed transposase adapters with Trim_Galore with a minimum length of 20 in paired-end mode. We aligned trimmed reads using Bowtie2 (Langmead and Salzberg, 2012) with default parameters. The Bowtie2 index was constructed with the default parameters for the GRCh38 reference genome. We filtered out reads that mapped to chrM and used SAMtools (Li et al., 2009b) to filter out reads with MAPQ < 10. Additionally, duplicate reads were discarded using Picard Chromatin accessibility peaks were identified with MACS3 (Zhang et al., 2008) under default parameters and ‘--nomodel --nolambda -- keep-dup all --call-summits’. The count of absolute peaks per cell type refers to the number of peak regions reported in the ‘narrowPeak’ file (peaks with multiple summits are only counted once). The peak count estimates were adjusted by sample read depth.

### *In silico* annotation for isoforms

For each isoform, we annotated whether it encodes a protein, causes nonsense-mediated decay (NMD) (Tardaguila et al., 2018), contains repetitive elements (Smit et al.), is a transmembrane protein (Krogh et al., 2001), contains a domain motif (Mistry et al., 2021), contains a signal peptide (Almagro Armenteros et al., 2019), or has intrinsically disordered regions (IDRs) (Dosztányi, 2018) *in silico* based on the nucleotide sequence obtained by long-read sequencing. The Kozak context sequence was scored as Gcc[AG]ccatgG. The most important nucleotides are +4, −3, and −6, and they were scored as +3 and the others as +1 if they were matched (the maximum score is 13). To annotate inserted repetitive elements, we used the isoform sequence through the gene body and the reference genome (GRCh38) sequence for upstream and downstream (±1 kb) of the gene body.

### Proteome analysis by nanoLC/MS/MS

We referred to ongoing proteome analysis data by liquid chromatography (LC)/mass spectrometry (MS)/MS-based global proteomics and protein terminomics (Nishida and Ishihama et al., manuscript in preparation) to validate whether predicted ORFs encoded in isoforms in the Immune Isoform Atlas were translated. For sample preparation, 1 × 10^7^ of THP-1 cells (ATCC; American Type Culture Collection) with/without 10 ng/ml of Phorbol 12-myristate 13-acetate (PMA) treatment for 72 h and LCL cells (NA12878, Coriell Institute) with/without 50 ng/ml IFN-α2 for 6 h were lysed with phase transfer surfactant buffer (Masuda et al., 2008) followed by digestion with Lys-C/trypsin. These four samples were divided into four fractions each (16 samples in total) and labeled with 16-plexed TMTpro reagents to prepare a single TMT set. For protein terminomics, the TMT set was used to isolate protein terminal peptides using a strong cation exchange (SCX) chromatography system consisting of an Agilent 1260 Infinity II Bio-Inert LC with a BioIEX SCX column (250 mm × 4.6 mm, 5 μm, nonporous) (Santa Clara, CA), as described previously (Tsumagari et al., 2021). The isolated peptides were fractionated into 24 vials by reversed-phase HPLC at high pH conditions using a Nexera X2 system (Shimadzu, Japan, Kyoto) with a L-column 3 (2.1 mm × 150 mm, 3 μm, 110 Å). We also conducted protein C-terminomics using the CHAMP protocol (Nishida and Ishihama, 2022). In brief, the cell lysates were digested by V8 protease. After dividing each sample into four fractions, the protein C-terminal peptides were isolated using CeO_2_ chromatography. The 16 multiplexed samples were mixed to prepare a single TMT set and fractionated into 24 vials by reversed-phase HPLC as described above. For global proteomics, the tryptic digests of four different samples were divided into four fractions each (16 samples in total) and labeled with 16-plexed TMTpro reagents to prepare a single TMT set and fractionated into 24 vials by reversed-phase HPLC as described above. All samples were desalted by SDB-StageTips (Rappsilber et al., 2007). NanoLC/MS/MS measurement was performed on an Orbitrap Exploris 480 mass spectrometer (Thermo Fisher Scientific, Waltham, MA) and an Ultimate 3000 LC system with a self-pulled needle column (250 mm, 100 μm ID) packed with Reprosil-Pur 120 C18-AQ 1.9 μm (Dr. Maisch, Ammerbuch, Germany). The flow rate was 400 nL/min. The LC mobile phases consisted of solvent A (0.5% acetic acid) and solvent B (0.5% acetic acid and 80% acetonitrile). The gradient was set as follows: 5–10% B in 2.5 min, 10–19% B in 57.8 min, 19–29% B in 21 min, 29–40% B in 8.7 min, and 40– 99% B in 0.1 min, followed by 99% B for 5 min. The electrospray voltage was set to 2.4 kV in the positive mode. For the survey scan, the mass range was from 375 to 1600 *m/z* with a resolution of 60,000, 100% normalized AGC target, and auto maximum injection time. For the MS/MS scan, the first mass was set to 110 *m/z* with a resolution of 45,000, 0.7 *m/z* of isolation window, 100% normalized AGC target, and auto maximum injection time. Fragmentation was performed by higher-energy collisional dissociation with a normalized collision energy of 30. The dynamic exclusion time was set to 60 s.

### Proteome data analysis

The MS raw files were searched to identify peptides by MaxQuant. To identify the novel isoforms, we customized the database. For LCL, we quantified isoforms by remapping the RNA-seq dataset of the GEUVADIS (Genetic European Variation in Disease) project (Lappalainen et al., 2013) (n = 463, EMBL-EBI, E-GEUV-1). For THP-1, we quantified isoforms by remapping the RNA-seq dataset derived from naïve THP-1 cells (n = 3, deposited in the GEO under the GSE157052). Considering sample size, we filtered out isoforms with minimum TPM for all samples = 0 and < 2 for LCL and THP-1, respectively. In addition, to predict novel proteins, we deleted isoforms whose entire predicted ORFs were included in GENCODE. As a result, predicted ORFs encoded by 16,190 isoforms were retrieved as novel isoform candidates. Then, we constructed a non-redundant protein database by combining them with the Swiss-Prot database of human proteins including isoforms (42,360 entries, 2022_06) for the database search in this study.

For tryptic peptides, methionine oxidation and protein N-terminal acetylation were selected as variable modifications, and cysteine carbamidomethylation and peptide N-terminal and lysine TMTpro labels as fixed modifications. For V8 protease-digested peptides, methionine oxidation was selected as a variable modification and cysteine carbamidomethylation and peptide N-terminal and lysine TMTpro labels as fixed modifications. A maximum of two missed cleavages were allowed. The FDR filter was set to 1% at both the peptide-spectrum match (PSM) and protein levels. MS/MS spectra were manually inspected, and the acceptance criterion was set at an andromeda score > 80, which corresponds to an FDR of < 0.075% at the PSM level.

### Short-read RNA sequencing and processing

We utilized the datasets (single nucleotide polymorphism (SNP) array and RNA-seq data) of previous expression quantitative trait locus (eQTL) studies obtained from four Europeans cohorts: the EvoImmunoPop project (Quach et al., 2016) (European Genome-phenome Archive [EGA], EGAS00001001895), the DICE (database of immune cell expression, expression quantitative trait loci, and epigenomics) project (Schmiedel et al., 2018) (the database of Genotypes and Phenotypes (dbGaP), phs001703.v1.p1), the Immune Variation (ImmVar) study (Lee et al., 2014), (dbGaP, phs000815.v1.p1), and the GEUVADIS project (Lappalainen et al., 2013) (EMBL-EBI, E-GEUV-1).

We additionally performed genotype imputation using SNP array data. Pre-imputation quality control (QC) of the genotyping data was performed using PLINK (Purcell et al., 2007) with the following parameters (*--mind 0.02 --king-cutoff 0.0884 --geno 0.01 --maf 0.01 --hwe 1e-5*). Post-QC variants were prephased using SHAPEIT (Delaneau et al., 2012), and imputation was performed using MiniMac3 (Das et al., 2016) and 1000 Genomes Phase 3 (release 5) as the reference panel (Auton et al., 2015). Post-imputation QC was performed using PLINK with the following parameters (*--minimac3-r2-filter 0.3*). Genotyped and imputed SNPs or indels with minor allele frequency (MAF) ≥ 0.01 were used for subsequent QTL analysis with related expression datasets.

For RNA-seq, 3′ ends with low-quality bases (Phred quality score < 20), and adaptor sequences were trimmed using Trim_Galore from sequenced reads. We realigned the trimmed reads on the GRCh38 genome using STAR (Dobin et al., 2013) in two-pass mode with the *de novo* transcript annotations derived from long-read RNA-seq of 29 immune cell types.

Expression was quantified using StringTie2 (Pertea et al., 2015) and kallisto (Bray et al., 2016) independently using generated bam files from STAR and trimmed reads, respectively. For gene-level quantification, we combined all isoforms of a gene into a single transcript as described elsewhere (Aguet et al., 2017). Raw read counts were normalized with the Transcripts Per Kilobase Million (TPM) method (Li et al., 2009).

### Cell-type-specific isoforms

After filtering out isoforms with low expression levels (reads per million, RPM > 2) and isoform ratios in each related gene (isoform ratio > 0.2), cell-type-specific isoforms were identified based on their Shannon entropies using the ROKU (Kadota et al., 2006) function in the TCC package (Sun et al., 2013).

### RBP binding analysis

To investigate the association between RBPs and specifically expressed isoforms, we searched RBP binding motifs in the sequence of the 3′-UTR of each isoform using RBPmap (Paz et al., 2014). Then, the numbers of predicted binding sites for each RBP by region (5′-UTR, ORF, and 3′-UTR) were aggregated and compared between specifically expressed isoforms and others.

### Translational efficiency

We utilized RNA-seq and Ribo-seq datasets obtained from 52 common Yoruba individuals among the RNA-seq dataset derived from the GEUVADIS project (Lappalainen et al., 2013) (EMBL-EBI, E-GEUV-1) and the Ribo-seq dataset deposited in GEO under GSE61742 (Battle et al., 2015), respectively. To calculate translational efficiency at the isoform level, trimmed reads were aligned to the *de novo* transcriptome sequences generated from long-read sequencing for 29 immune cell subsets using STAR (Dobin et al., 2013) as with tools developed for the same purpose (Reixachs-Solé et al., 2020; Wang et al., 2016). Then, we applied generated bam files to the *coverageDepth* and *translationalEfficiency* functions with corrections using the maximum translational efficiency value in the 90 most highly ribosome-occupied nucleotides window within the feature in *ribosomeProfilingQC* R package (Ingolia et al., 2009; Ou J and Hoye M, 2022). We calculated the translational efficiency only of isoforms that satisfied coverage depth > 1 of both Ribo-seq and RNA-seq (86967 isoforms) to avoid the potential over-estimating of translational efficiency due to low coverage.

### Secondary structure

To estimate the presence of local RNA secondary structures, a window length of 25 nucleotides was moved at the step size of one nucleotide, starting from TSS to TTS, and the Gibbs free energy (Δ*G*) was calculated as the predicted local folding strength for each window using the RNAfold program (Gruber et al., 2008). Then, we tested the correlation translational efficiency of a particular isoform and Δ*G* values at each scaled position relative to the TSS (0 = TSS, 1, 2, …, 100 = first nucleotide of the ORF, 101, …, 200 = last nucleotide of the ORF, 201, …, 300 = TTS), and correlation coefficients were averaged over isoforms at each position.

To validate predicted local folding strength, we tested correlations with the Parallel Analysis of RNA structure (PARS) score (Kertesz et al., 2010; Wan et al., 2014) (**Figure S4**). To calculate PARS scores, we downloaded RNA fragments generated from LCLs of a family trio with treatment by RNase V1 or S1 nuclease deposited in GEO under GSE50676 (Wan et al., 2014) and mapped on the GRCh38 genome using STAR (Dobin et al., 2013) in two-pass mode with the *de novo* transcript annotations derived from long-read sequencing for 29 immune cell types. Then, we quantified the number of double-stranded reads (V1) and single-stranded reads (S1) that were initiated on each base on the RNA. The read counts of double and single stranded reads for each sequencing sample were normalized by sequencing depth. For a particular isoform with *N* bases in total, the PARS score of its *i*th base was defined by the following formula where V1 and S1 are normalized V1 and S1 scores, respectively. A small number (5) was added to reduce the potential over-estimating of structural signals of bases with low coverage:

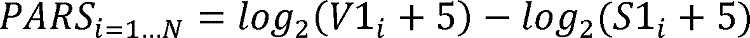

After removing the scaled position with all PARS scores being zero among three samples, we tested the correlation between averaged PARS score and predicted local folding strength at each scaled position. We capped the PARS score to ± 7. We calculated PARS scores of only isoforms whose median TPM were the highest in each gene to minimize noise from multi-mapping. We quantified isoform expression values using LCL samples in the GEUVADIS cohort (Lappalainen et al., 2013). In total, PARS scores of 5800 isoforms were calculated.

### Isoform-switch analysis

We downloaded short-read RNA-seq datasets obtained from clinical patients and healthy controls from the GEO through accession numbers GSE72509 for SLE and GSE89408 and GSE118829 for RA. From raw sequenced reads, 3′ ends with low-quality bases (Phred quality score < 20) and adaptor sequences were trimmed using Trim_Galore. An expression matrix was generated using kallisto (Bray et al., 2016) with default parameters and the *de novo* transcript annotations derived from long-read sequencing for 29 immune cell types. Then, we used IsoformSwitchAnalyzeR in the R package to detect genes that have changed splicing patterns (Vitting-Seerup et al., 2019). Briefly, IsoformSwitchAnalyzeR measures isoform usage via n isoform fraction (IF) values, which quantifies the fraction of the parent gene expression originating from a specific isoform (calculated as isoform_exp / gene_exp). Consequently, the difference in isoform usage is quantified as the difference in isoform fraction (dIF) calculated as IF2 − IF1, and these dIF are used to measure the effect size (like fold changes are in gene/isoform expression analysis). The cutoff for dIF and FDR to detect significantly switched isoforms were 0.05 and 0.05, respectively.

### QTL analysis

Independent QTL analysis was performed for each condition. In common with all QTL analyses, the following process was performed: 1) normalization of the expression matrix was performed with quantile normalization, rank-transformed normalization, and PEER normalization using 15 hidden factors for all QTL analyses (Stegle et al., 2012), and 2) the variants with MAF ≥ 0.01 within a 1-megabase (Mb) window around each transcript using MatrixEQTL of the R package (Shabalin, 2012) with the top 10 genetic principal components as covariates.

The pipeline for sQTL analysis was the same as in gene-level eQTL analysis, except for the preparation of the expression matrix. A comparison of two representative quantification methods, 1) alignment-based transcript quantification and 2) alignment-free transcript quantification, showed a high correlation in expression values at the gene level, but a moderate correlation at the isoform level between the two methods (**Figure S7**). Therefore, we utilized the independent isoform ratio in related genes derived from StringTie2 (Pertea et al., 2015) and kallisto (Bray et al., 2016), which are alignment-based transcript quantifications and alignment-free transcript quantifications for trQTL analysis, respectively. In addition, to capture differences in coding-sequence more sensitively, we used the isoform ratio quantified by kallisto after clustering the isoforms with completely identical coding-sequences by VSEARCH (Rognes et al., 2016). We also conducted junction-based sQTL analysis using LeafCutter (Li et al., 2018).

With regard to 3′aQTL, we used dynamic analyses of APA from the RNA-seq (DaPars) algorithm (Masamha et al., 2014; Xia et al., 2014) to identify APA events. The multi-sample DaPars v.2 regression framework calculates the percentage of the distal poly(A) site usage index (PDUI) value for each gene in each condition. Subsequently, we analyzed the association between variants within 1-Mb from the 3′-UTR region and quantile- and rank-normalized PDUI values with covariates, the same as with eQTL and sQTL analyses. To investigate the enriched function of genes with 3′aQTL effect, pathway analysis was performed using Enrichr (Chen et al., 2013).

### Stratified linkage disequilibrium score regression analysis (S-LDSC)

We extracted eQTL, sQTL, and 3′aQTL variants (SNPs or indels with nominal p-values < 1 × 10^-5^) and performed S-LDSC (Finucane et al., 2015) adjusting for functional annotation (baseline model v1.2 provided by the developers). Formatted GWAS summary statistics for S-LDSC by developers were downloaded from https://alkesgroup.broadinstitute.org/sumstats_formatted/.

### Colocalization analysis of QTL and GWAS

To evaluate the colocalization of QTL and GWAS signals, we applied a Bayesian framework using coloc of the R package (Giambartolomei et al., 2014). We tested for a 500,000 bp window centered on the GWAS lead variant and considered PP-H4 (posterior probability of shared causal variant) > 0.8 as a significant colocalization. Formatted GWAS summary statistics for S-LDSC by developers were downloaded from https://alkesgroup.broadinstitute.org/sumstats_formatted/ and severe COVID-19 from the COVID-19 HGI release5 (https://storage.googleapis.com/covid19-hg-public/20201215/results/20210107/COVID19_HGI_A2_ALL_eur_leave_23andme_20210107.b37.txt.gz).

### Statistical test

The statistical tests performed are indicated in the figure legends or STAR Methods.

### Supplementary Figure legends

**Figure S1.**
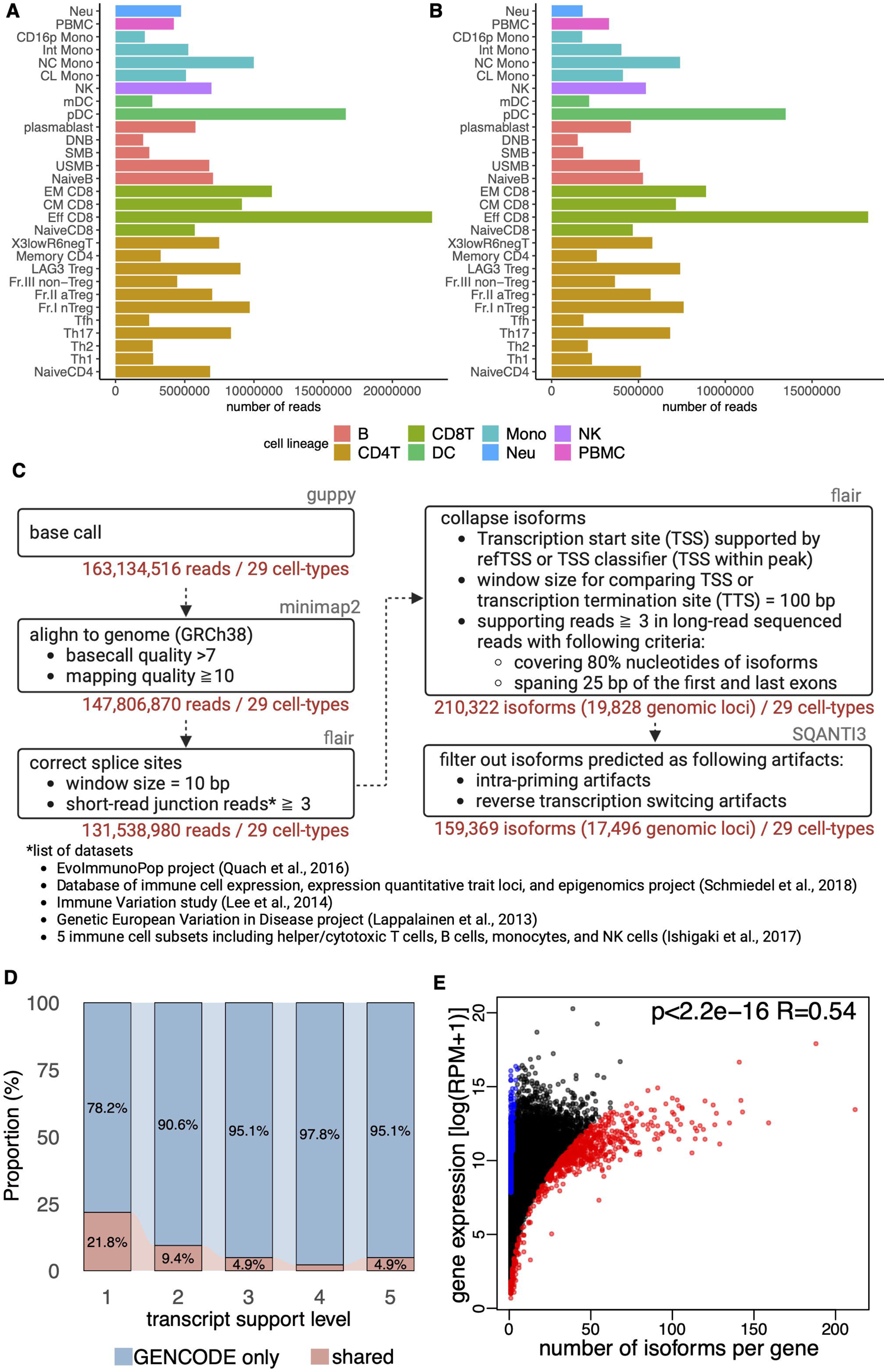
Workflow for generating Immune Isoform Atlas, related to Figure 1. (A) Raw sequenced reads in each cell subset. (B) Mapped sequenced reads on the genome in each cell subset after filtering out reads with a mapping quality (MAPQ) of 10 or less. (C) Workflow for processing long-read sequenced reads to generate the Immune Isoform Atlas. The gray letters in the upper right corner of the box indicate the name of the tool for each process. The red letters in the lower right corner indicate the number of reads or isoforms that passed the process. (D) The proportion of isoforms that are shared between the Immune Isoform Atlas and GENCODE (red) and registered only in GENCODE (blue) according to transcript support levels. In GENCODE, isoforms are scored according to how well mRNA and EST alignments match over its full length as follows: 1, all splice junctions of the transcript are supported by at least one non-suspect mRNA; 2, the best supporting mRNA is flagged as suspect or the support is from multiple ESTs; 3, the only support is from a single EST; 4, the best supporting EST is flagged as suspect; 5, no single transcript supports the model structure. (E) Scatter plot of the number of alternatively spliced isoforms per gene and related gene expression. The red and blue dots indicate genes with the top 5% and bottom 5% of the number of alternatively spliced of isoforms after correction by expression levels of related genes, respectively. The expression value was normalized by reads per million (RPM).

**Figure S2.**
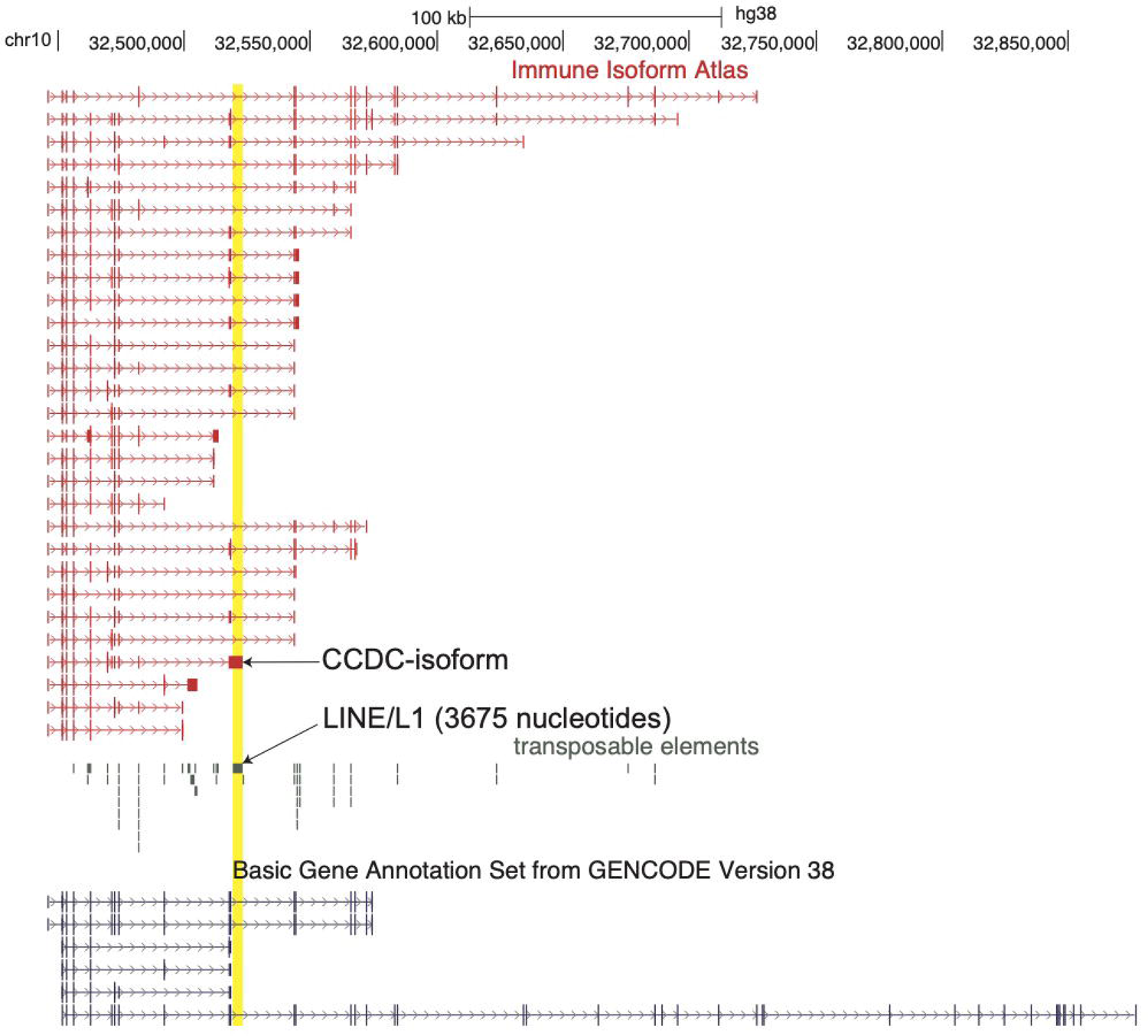
Example of isoform inserted transposable elements, related to Figure 3. These isoforms were transcribed from the *CCDC7* locus, and a particular isoform had an inserted *LINE/L1* highlighted in yellow in the 3′-UTR. The inserted 3′-UTR was not registered in GENCODE, as shown at the bottom.

**Figure S3.**
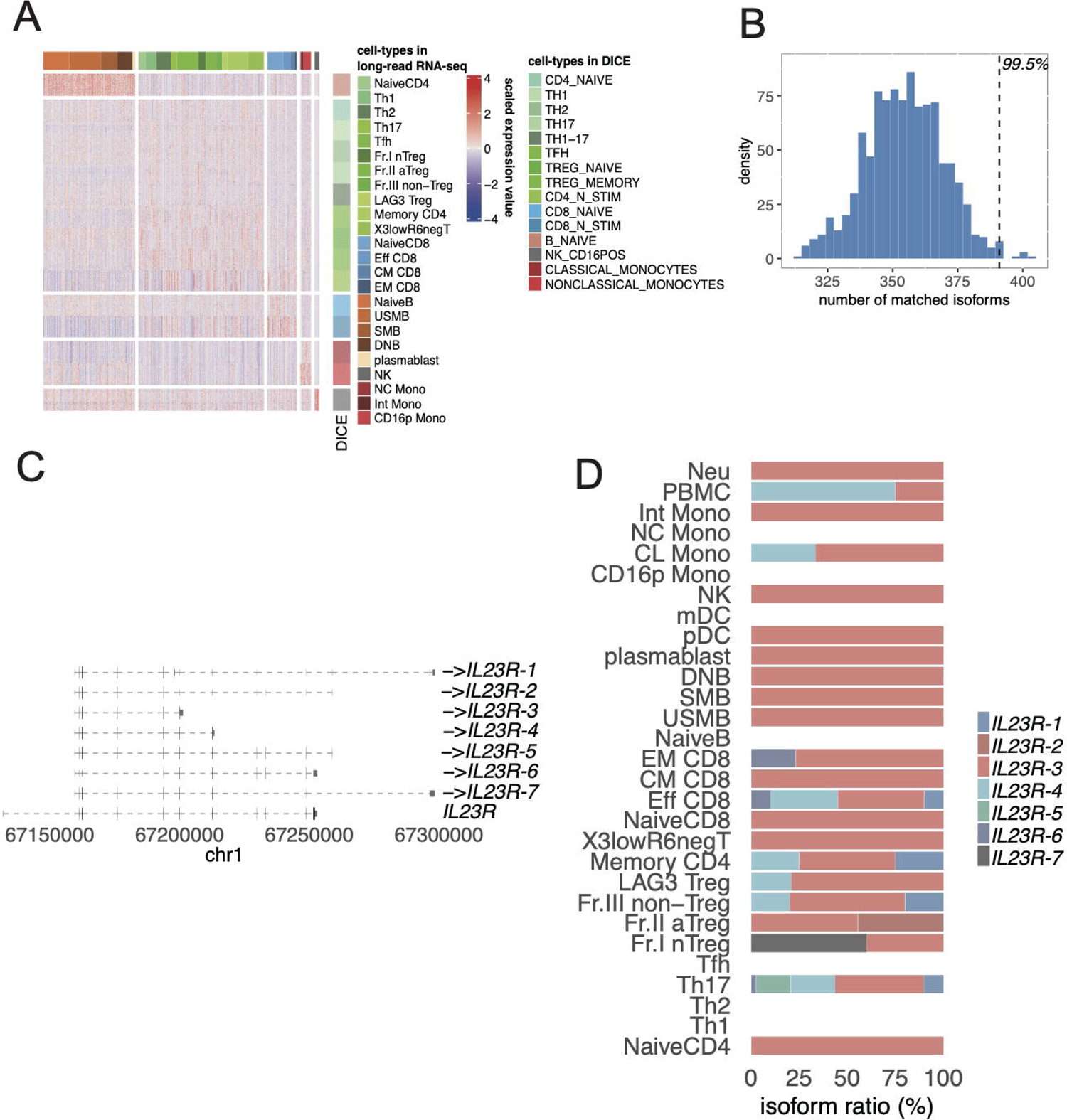
The concordant expression of cell-type-specific isoforms in relevant cell types of short-read RNA-seq datasets, related to Figure 4. (A) Expression of cell-type-specific isoforms (Figure 4B) in cell subsets of short-read RNA-seq datasets (Schmiedel et al., 2018). Column-wise Z scores of normalized counts are plotted. Cell types are arranged by hierarchical clustering. The colored bars indicate the cell types in each dataset. (B) Histogram of the number of matched counts between the labels of specifically expressed cell types in the long-read sequencing dataset and the label of the highest expressed cell type in the short-read sequencing dataset by the 1,000 permutation tests. The vertical line shows the true counts of matched isoforms (395) between them (permutation p-value = 0.005). (C) Example of the specifically expressed isoform transcribed from *IL23R* locus. The collapsed gene structure registered in GENCODE is shown at the bottom. The arrow indicates the direction of transcription in the genome. (D) Isoform ratio in *IL23R*.

**Figure S4.**
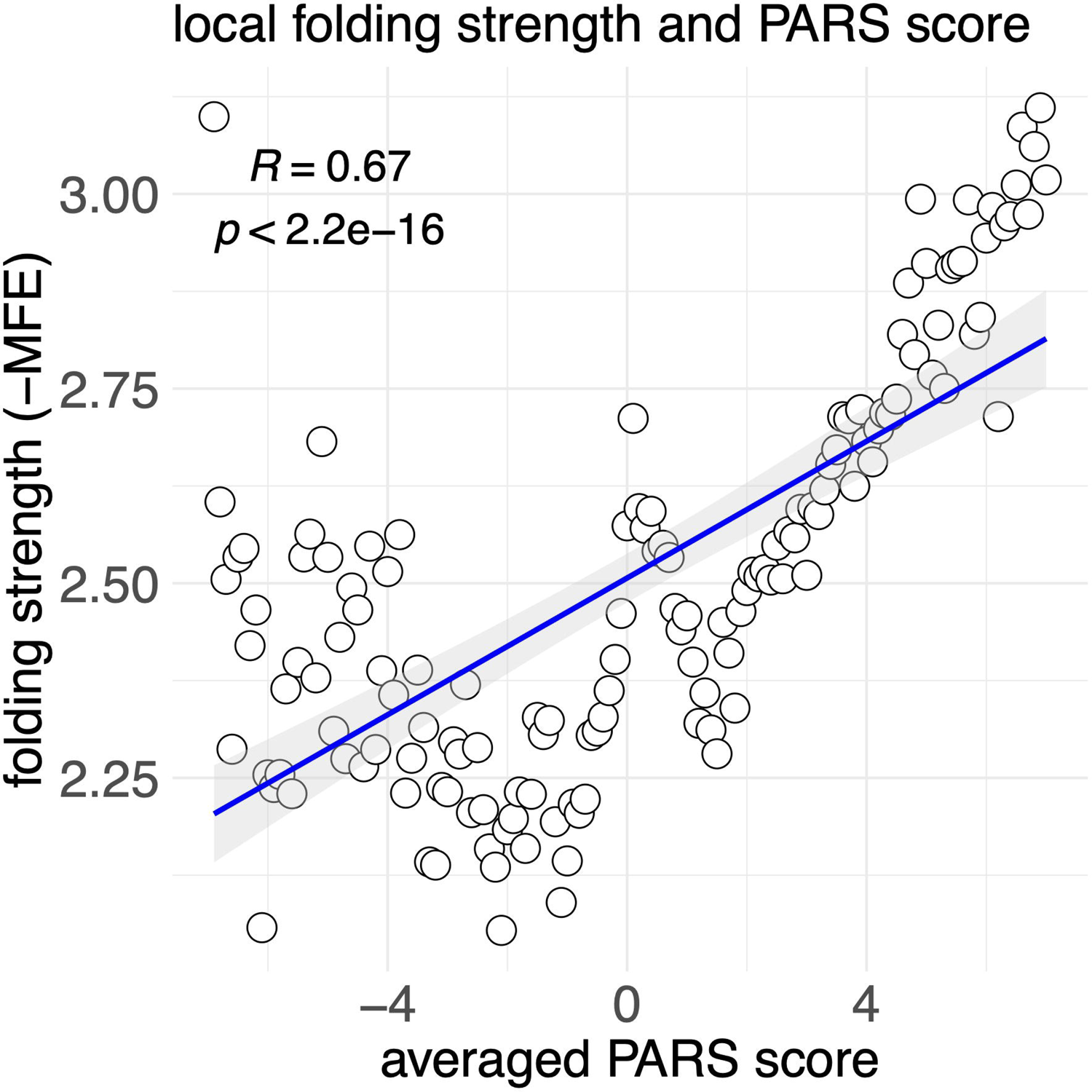
Correlation between Parallel Analysis of RNA structure (PARS) scores and predicted local folding strength of isoforms (Methods), related to Figure 5.

**Figure S5.**
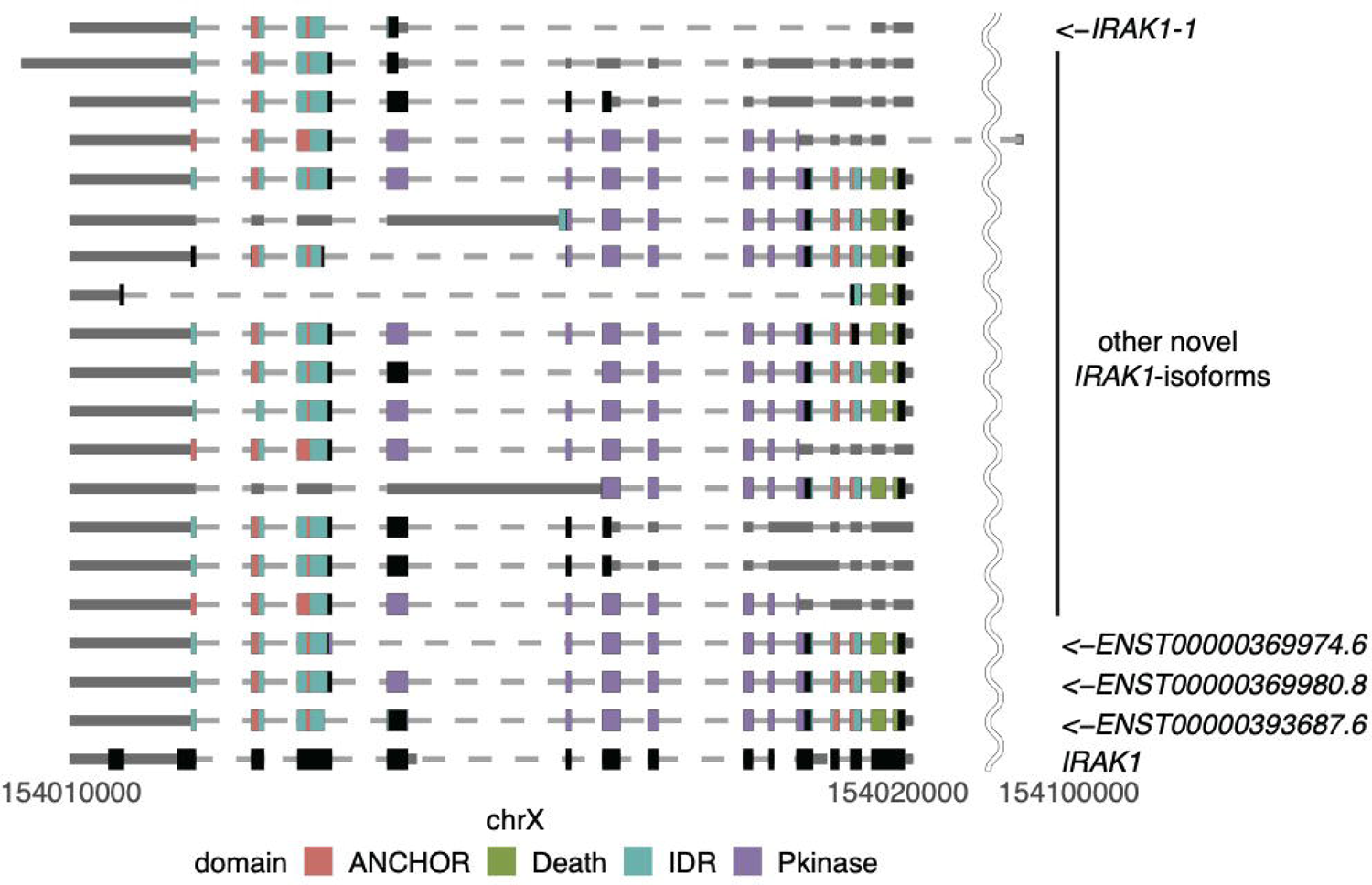
The structure of all isoforms that are transcribed from the *IRAK1* locus, related to Figure 6. The collapsed gene structure registered in GENCODE is shown at the bottom. The arrow indicates the direction of transcription in the genome.

**Figure S6.**
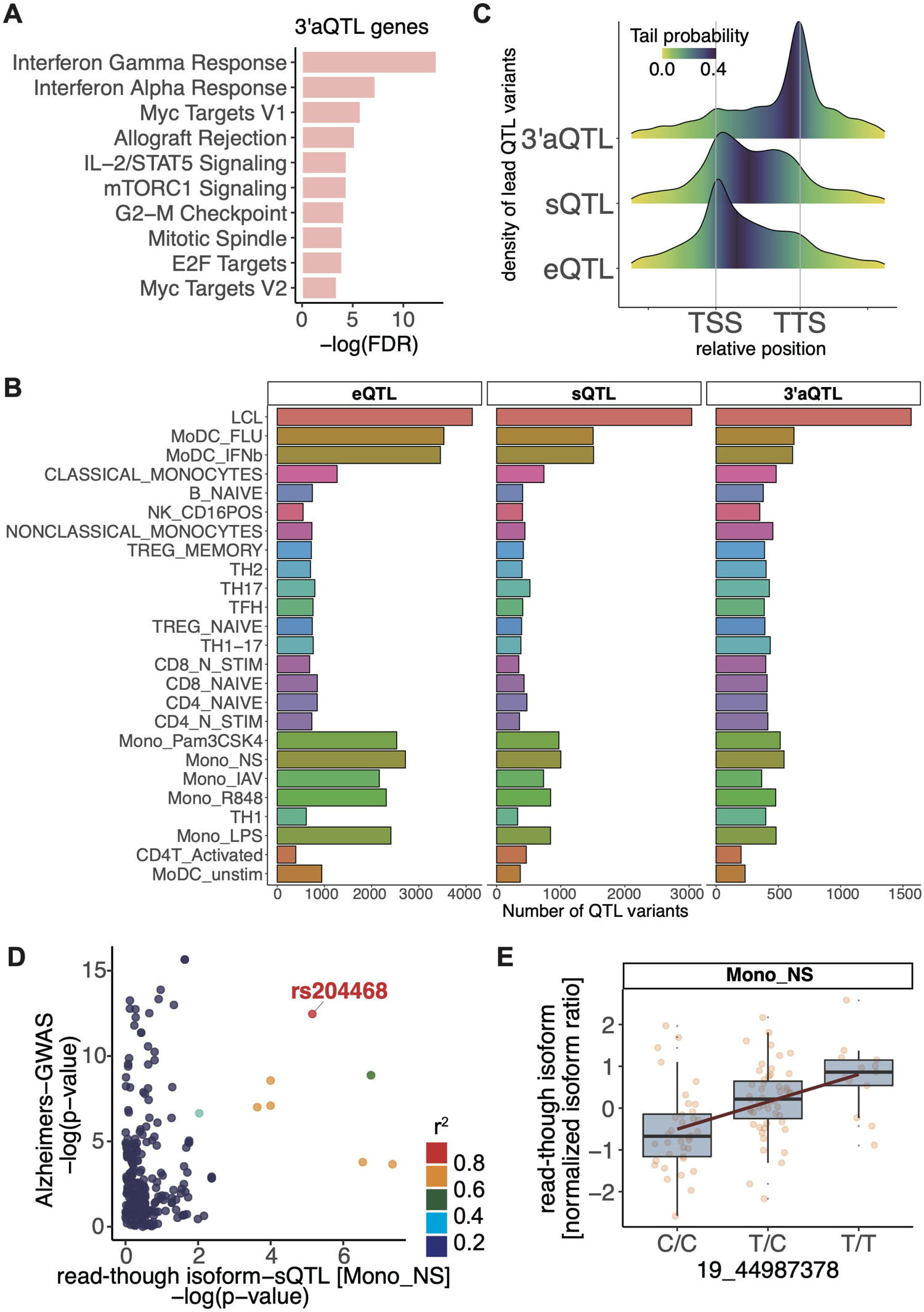
Characteristics of QTL genes and QTL variants, related to Figure 7. (A) Characteristics of alternative polyadenylation (APA) quantitative trait loci (3′aQTL)-genes. Enriched pathways in 3′aQTL-genes (B) The number of total lead QTL variants (nominally significant p-values < 1 × 10^-5^) for each eGene (eQTL), junction (sQTL), and isoform (3’aQTL), sorted by the sample sizes. (C) The distribution of lead QTL variants in the relative position. (D) Colocalization plot of trQTL of the read-through isoform transcribed from the *TOMM40_APOE* locus and GWAS of Alzheimer’s disease in non-stimulated monocytes. (E) QTL plot of normalized isoform ratio of the read-through isoform transcribed from the *TOMM40_APOE* locus in non-stimulated according to each genotype of rs204468. Abbreviations of cell-types in short-read RNA-seq datasets and their related cohorts were as follows: LCL, lymphoblastoid cell-lines [GEUVADIS]; Mono_NS, monocytes without treatment [EvoImmunoPop]; Mono_LPS, monocytes treated with TLR4 agonist [EvoImmunoPop]; Mono_Pam3CSK4, monocytes treated with TLR 1/2 agonist [EvoImmunoPop]; Mono_R848, monocytes treated with TLR 7/8 agonist [EvoImmunoPop]; Mono_IAV, monocytes treated with H1N1 Influenza A virus [EvoImmunoPop]; CD4T_Activated, CD4^+^ T cells treated with anti-CD3/CD28 antibody [ImmVar]; MoDC_unstim, monocyte-derived dendric cells without treatment [ImmVar]; MoDC_IFNb, monocyte-derived dendric cells treated with IFNb [ImmVar]; MoDC_FLU, monocyte-derived dendric cells treated with H1N1 Influenza A virus [ImmVar]; CD4_NAIVE, naïve CD4^+^ T cells [DICE]; CD4_N_STIM, naïve CD4^+^ T cells treated with anti-CD3/CD28 antibody [DICE]; TH1, T_H_1 cells [DICE]; TH2, T_H_2 cells [DICE]; TH1-17, T_H_1-17 cells [DICE]; TH17, T_H_17 cells [DICE]; TFH, T_FH_ cells [DICE]; TREG_NAIVE, naïve T_REG_ cells [DICE]; TREG_MEMORY, memory T_REG_ cells [DICE]; CD8_NAIVE, naïve CD8^+^ T cells [DICE]; CD8_N_STIM, Naïve CD8^+^ T cells treated with anti-CD3/CD28 antibody [DICE]; B_NAIVE, naïve B cells [DICE]; CLASSICAL_MONOCYTES, classical monocytes [DICE]; NONCLASSICAL_MONOCYTES, nonclassical monocytes [DICE]; NK_CD16POS, NK cells [DICE].

**Figure S7.**
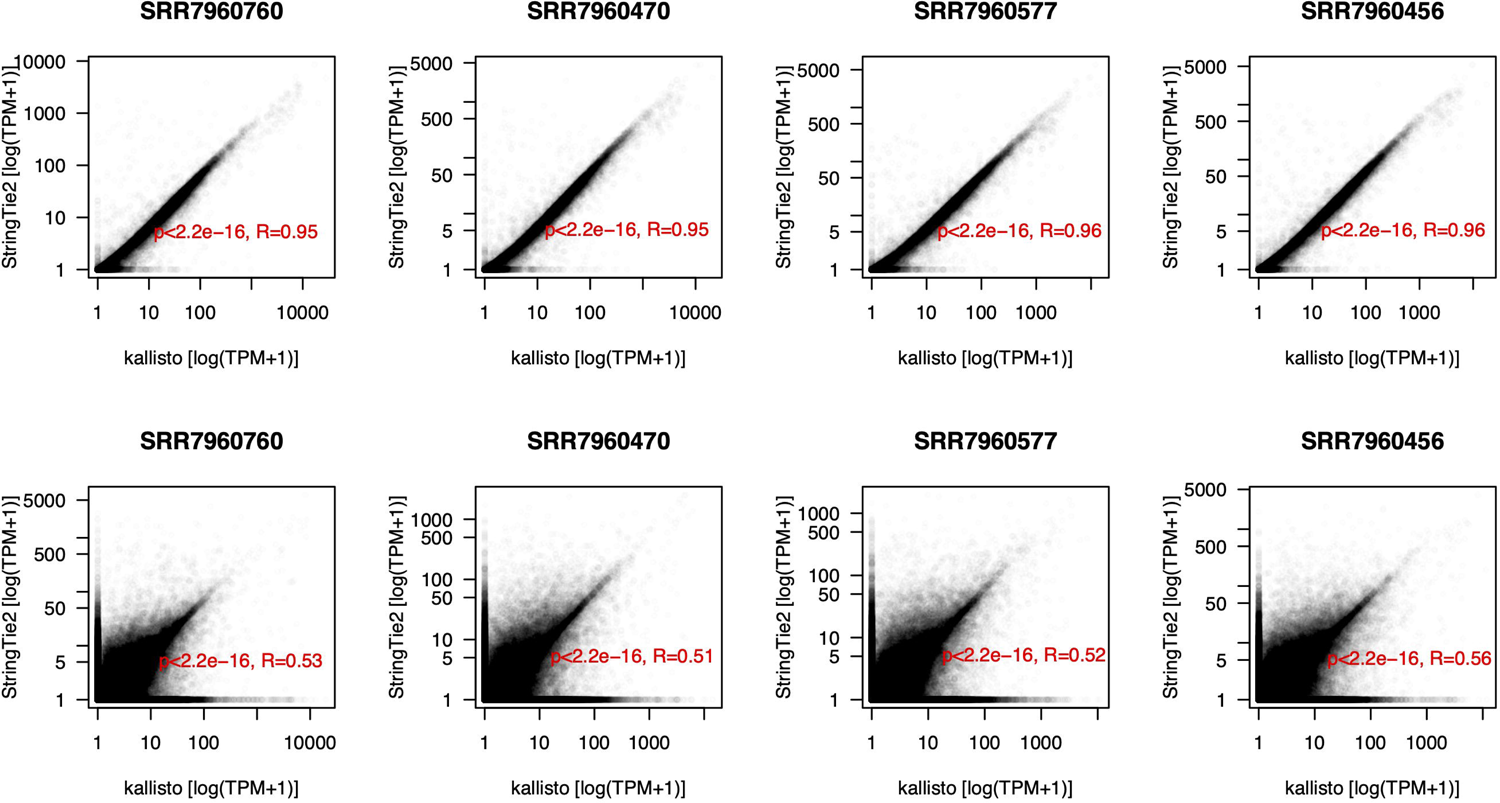
Comparison of quantification tools, related to STAR Methods. The top four and bottom four scatter plots were drawn using expression values at the gene levels and isoform level, respectively. Each dot represented genes (top) and isoforms (bottom). Four samples were randomly selected from short-read RNA-seq datasets of DICE. The boxes contain Spearman’s correlation coefficient and p-value.

## Notes

### Competing Interest Statement

The authors have declared no competing interest.

